# Design of an exclusive obligate symbiosis for organism-based biological containment

**DOI:** 10.1101/2025.01.05.631398

**Authors:** Amanda M. Forti, Michaela A. Jones, Defne N. Elbeyli, Neil D. Butler, Aditya M. Kunjapur

## Abstract

Biological containment is a critical safeguard for genetically engineered microbes prior to their environmental release to prevent proliferation in unintended regions. However, few biocontainment strategies can support the longer-term microbial survival that may be desired in a target environment without repeated human intervention. Here, we introduce the concept of an orthogonal obligate commensalism for the autonomous creation of environments that are permissive for survival of a biocontained microbe. We engineer one microbe to produce a non-standard amino acid (nsAA), and we engineer synthetic auxotrophy in a second microbe via reliance on this nsAA for growth. We show that this obligate commensalism is highly effective, with the survival of our commensal organism during co-culture dependent on the presence of our producer strain. We also show that this commensalism is orthogonal to a small microbial consortium isolated from maize roots, with survival of the synthetic auxotroph conditional upon the presence of the nsAA-producing strain in the consortium. Overall, our study demonstrates a transition from a chemical to a biological dependence for biocontained organisms that could lay the groundwork for biocontained synthetic ecologies.

## Introduction

Synthetic biology offers the promise of addressing issues in agriculture, environmental health, and public health by direct release of genetically engineered microbes into target environments^1–3^. However, a grand challenge that has limited the use of engineered microbes in the environment is the design of effective safeguards that provide strict spatiotemporal control over the survival of engineered organisms^3–5^. Because most naturally occurring auxotrophy can be overcome through microbial cross-feeding^5–9^, more sophisticated biological containment strategies are required to be effective in open systems. The last decade has featured the demonstration of several innovative intrinsic biocontainment strategies, including genetic circuits that trigger cell death in response to specific molecules or environmental conditions^10–14^ as well as synthetic auxotrophy, where the microbe requires a synthetic nutrient for its survival, most commonly a non-standard amino acid (nsAA) whose incorporation is required for full-length translation or proper folding of essential proteins^15–19^. Synthetic auxotrophy has shown promise as an intrinsic biocontainment strategy given the low rates of escape from biocontainment observed, which includes bacterial strains that have not exhibited detectable escape even after months of continuous culturing^20^. These escape frequencies can be well below the NIH standard of 10^-8^ escapees per colony forming unit (escapees/CFU)^21^, though a different standard may be necessary for eventual environmental release.

While the demonstration that synthetic auxotrophs remain biocontained in monoculture is promising, further innovation is required for the implementation of biocontainment in the environment. In particular, desired applications ranging from bioremediation to plant growth promotion likely require survival of engineered microbes for timescales of weeks or months in target environments, preferably without repeated manual provision of synthetic nutrient (Fig. 1a). As such, the ability to create a sustained permissive condition in a target environment while maintaining a non-permissive condition in the broader environment poses a potential paradox. In our view, an attractive solution would transform the chemical dependence of a synthetic auxotroph (an nsAA “utilizer”) into a biological dependence on exclusively one other engineered organism (an nsAA “producer”). This unidirectional exclusive reliance should generate a sustained and local permissive environment for the utilizer microbe upon fixing the position of the producer organism.

**Figure 1.**
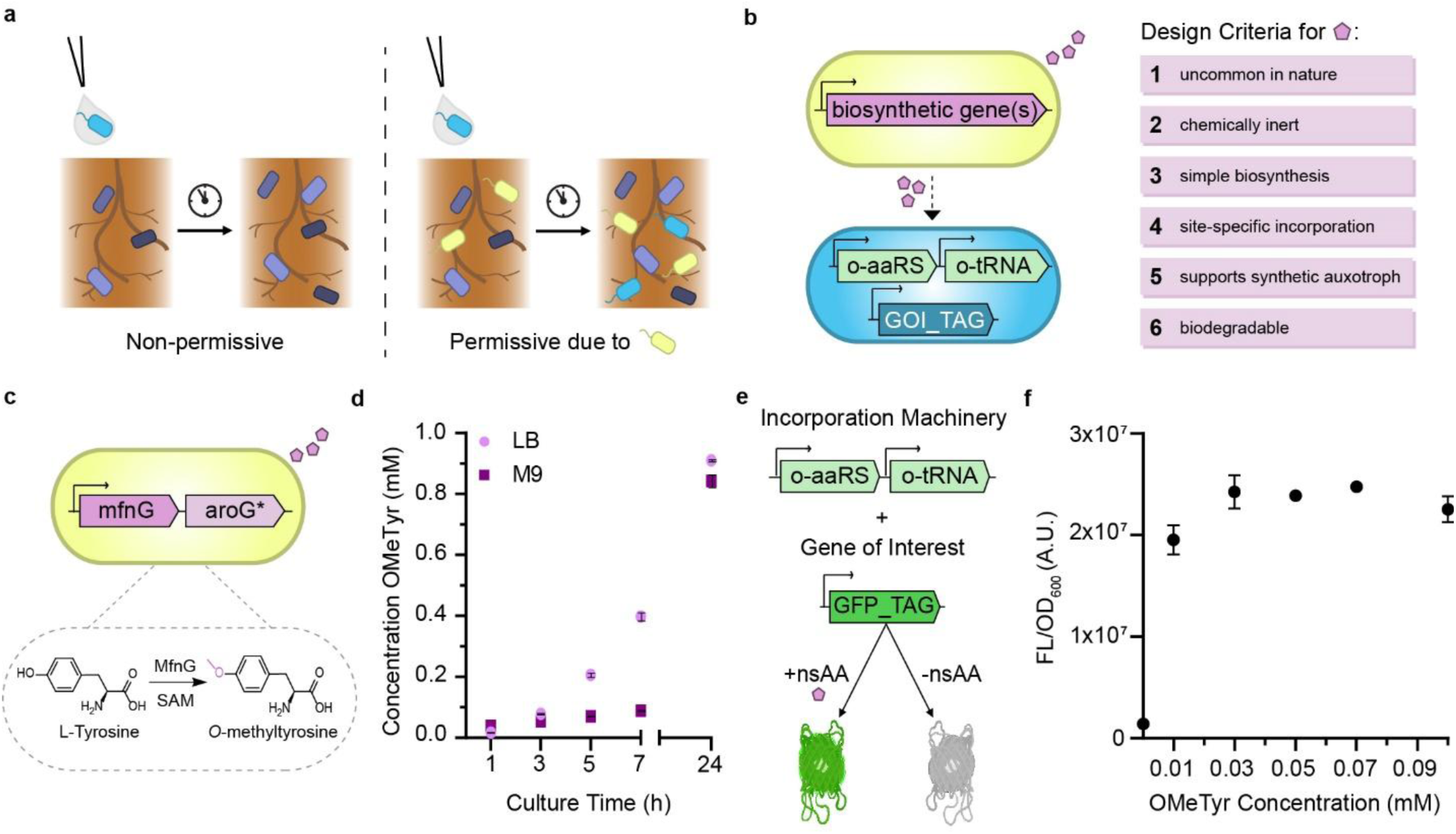
Characterization of the individual components envisioned to support an obligate commensal based on the non-standard amino acid OMeTyr. **a,** Illustration of the concept of an organism engineered to be an orthogonal and obligate commensal (blue microbe) that strictly relies on the inclusion of another engineered organism (yellow microbe) within a community. **b,** Schematic for one proposed strategy to create a form of commensalism mediated by the biosynthesis of a non-standard amino acid (nsAA) by one organism followed by the utilization of the nsAA by another organism that requires it. Included are design criteria for suitable nsAAs. **c,** Depiction of the biosynthesis of the nsAA chosen for this study, *O*-methyl-L-tyrosine (OMeTyr), and the associated genetic construct that expresses *mfnG* and *aroG\** genes within a synthetic operon. **d,** Titers of OMeTyr as a function of time produced by an engineered *E. coli* strain that co-expresses *mfnG* and *aroG\** genes under a constitutive promoter in LB media or M9 minimal media with glucose. **e,** Depiction of the generalized orthogonal translation machinery used to achieve site-specific incorporation of OMeTyr in *E. coli*: An engineered aminoacyl tRNA synthetase-tRNA pair (o-AARS/o-tRNA) derived from *M. jannaschii* coupled with a fluorescent reporter (GFP_TAG) that requires suppression of an in-frame amber stop codon for full-length translation. **f,** Fluorescence (ex: 485 nm, em: 525 nm) normalized by OD_600_ resulting from the incorporation of increasing concentrations of OMeTyr supplied to cultures of the genomically recoded RR3 (DE3) *E. coli* strain that harbors the NapARS synthetase in an aromatic amino acid dropout version of MOPS EZ rich media with glucose 9 h after inoculation. Sample sizes are n=3 using biological replicates (samples originated from the same glycerol stock). Data shown are mean ± standard deviation.

Here, we design and characterize a synthetic exclusive reliance – or, in mixed parlance of synthetic biology and ecology, an orthogonal and obligate commensalism - of one *Escherichia coli* strain on another (Fig. 1b). Our proof-of-principle demonstration using two *E. coli* strains accomplishes our primary objectives of generating no detectable survival of the utilizer strain when co-inoculated with a non-producer strain and generating survival of the utilizer strain when co-inoculated with a producer strain. Encouragingly, the exclusive reliance of the utilizer strain on the producer strain endures even in a small consortium of undomesticated soil microbes. We conclude by discussing two visions for eventual deployment that illustrate the critical role this designed interaction should have in enabling spatial containment of engineered microbes released into the environment without requiring repeated manual intervention nor physical confinement of all released microbes.

## Results

### nsAA biosynthesis and incorporation

To design an orthogonal and obligate commensalism, we established criteria for an nsAA that could mediate a metabolic interaction between two organisms. An ideal nsAA would be: (1) uncommon in nature, to lower the risk of natural cross-feeding; (2) chemically inert, to prevent unwanted reactivity; (3) biosynthetically accessible by some combination of known metabolic pathways, to enable its engineered biosynthesis; (4) amenable to site-specific incorporation within proteins using existing genetically encoded tools, to enable its utilization; (5) capable of supporting survival of a synthetic auxotroph, to force commensalism; and (6) biodegradable, to maintain strict reliance on the nsAA producer for survival and to avoid environmental pollution. Previously reported *E. coli* synthetic auxotrophs use orthogonal translation systems (OTSs) for the incorporation of a variety of different nsAAs, all of which have unclear biosynthetic routes from simple carbon sources^15–18^. Instead, we built a system based on the nsAA *O*-methyl-L-tyrosine (OMeTyr), which met the first four criteria and was one of the first nsAAs site-specifically incorporated into proteins^22^. OMeTyr biosynthesis in cells that were supplied tyrosine-containing media was recently reported, as was the site-specific incorporation of biosynthesized OMeTyr within proteins in multiple heterologous hosts^23^. However, for the design of an orthogonal obligate commensalism, *de novo* synthesis of OMeTyr from simple carbon sources must be achieved so that biosynthesis is guaranteed in any nutritionally complete environment.

To investigate OMeTyr biosynthesis, we engineered an *E. coli* strain for heterologous expression of an *O-*methyltransferase from the deep sea sediment-derived *Streptomyces drozdowiczii*, MfnG^24^. This enzyme catalyzes the transfer of a methyl group from the donor co-factor *S*-adenosyl-L-methionine (SAM) to the substrate L-tyrosine (Fig. 1c). We employed a well-established strategy to increase flux through the aromatic amino acid biosynthesis pathway: expression of a feedback-resistant DAHP synthase (AroG_D146N, or AroG*)^25,26^. Encouragingly, after we co-expressed the *aroG\** and *mfnG* genes and grew cells in M9 minimal media supplemented with glucose as a sole carbon source, we observed a titer of 0.229 ± 0.084 mM OMeTyr after 24 h (Supplementary Fig. 1a). We designed a single operon to contain all genes of the required enzymes (Fig. 1c) and compared the use of a constitutive promoter with an inducible promoter. We found that the use of a constitutive promoter increased the rate of OMeTyr biosynthesis in LB media, generating 0.398 ± 0.013 mM OMeTyr within the first 7 h after inoculation (Fig. 1d) compared to the use of the original, inducible promoter when induced at inoculation, which only produced 0.210 ± 0.005 mM OMeTyr by 7 h (Supplementary Fig. 1b). While OMeTyr biosynthesis was slower in M9 minimal media compared to LB, the cultures reached similar final concentrations of 0.843 ± 0.021 mM in M9 minimal media and 0.911 ± 0.004 mM in LB (Fig. 1d). Overall, these results showed that our engineered strains could achieve OMeTyr biosynthesis autonomously, meaning from simple carbon sources, in media lacking tyrosine, and without the need for an additional chemical inducer.

After optimizing OMeTyr biosynthesis, we focused on the first of two elements that would be required to create an OMeTyr-dependent *E. coli* strain: site-specific incorporation of OMeTyr within target proteins. We screened a library of previously reported OTSs comprised of engineered *Methanocaldococcus jannaschii* aminoacyl-tRNA synthetase and tRNA pairs using the widely accepted approach of suppression of an amber stop codon located within a superfolder GFP reporter (GFP_TAG)^27^ (Fig. 1e). Encouragingly, after measuring GFP fluorescence normalized by optical density (FL/OD_600_) we discovered that several OTSs could facilitate site-specific incorporation of OMeTyr when supplied at 1 mM, including many not previously shown to have activity on OMeTyr (Supplementary Fig. 2). We chose to investigate the dose-response of our best performing synthetase, NapARS^28^, which exhibited a 22-fold increase in OD_600_ normalized GFP fluorescence when supplied with 1 mM OMeTyr compared to a 0 mM OMeTyr control. Importantly, this synthetase also had the lowest normalized fluorescence in the 0 mM OMeTyr condition among all the synthetases tested, indicating the highest fidelity for OMeTyr incorporation. Upon supplying lower concentrations of OMeTyr, we observed a greater than 13-fold increase in normalized GFP fluorescence even with as little as 0.01 mM OMeTyr supplied compared to the 0 mM nsAA condition (Fig. 1f).

### Distribution of nsAA biosynthesis and incorporation functions across two strains

After constructing OMeTyr biosynthesis and incorporation technology in separate strains, we next monitored and optimized the dynamics of biosynthesis, exchange, and incorporation of OMeTyr within a co-culture. We constructed one OMeTyr “producer” strain that harbored the plasmid containing constitutively expressed biosynthetic genes and another OMeTyr “utilizer” strain that harbored the plasmid containing inducible orthogonal translation machinery and reporter genes, each with common antibiotic resistance genes. We measured fluorescence to evaluate successful incorporation of biosynthesized OMeTyr in the utilizer strain (Fig. 2a). As a negative control, we also used a “non-producer” strain that carried a plasmid that harbored our desired antibiotic resistance but not the *mfnG* gene. When we varied the inoculation ratio of producer:utilizer strains in our co-culture, we saw an increase in fluorescence over the course of culture growth that was indicative of OMeTyr incorporation within GFP. We observed a 13.5-fold increase in maximum fluorescence between the co-culture containing the OMeTyr producer strain compared to the co-culture containing the non-producer strain (Fig. 2b). We also found that increasing the inoculation ratio of utilizer to producer increased the rate of GFP production (Fig. 2b). To verify the fidelity of our OMeTyr biosynthesis and incorporation system distributed across two strains, we conducted a larger scale co-culture, purified the GFP produced in co-culture, and performed intact protein mass spectrometry. The observed mass was 37322 Da, which aligned with the expected mass for GFP containing OMeTyr (Fig. 2c). As additional verification we performed in-gel tryptic digestion and LC-MS/MS on the same GFP protein, where 99.7 % of the protein purified contained OMeTyr and only 0.3 % contained misincorporated L-tyrosine. Having demonstrated that producer and utilizer organisms could achieve robust and high-fidelity incorporation of an orthogonal building block within a co-culture, we next set out to create a strain that would rely on OMeTyr for its survival.

**Figure 2.**
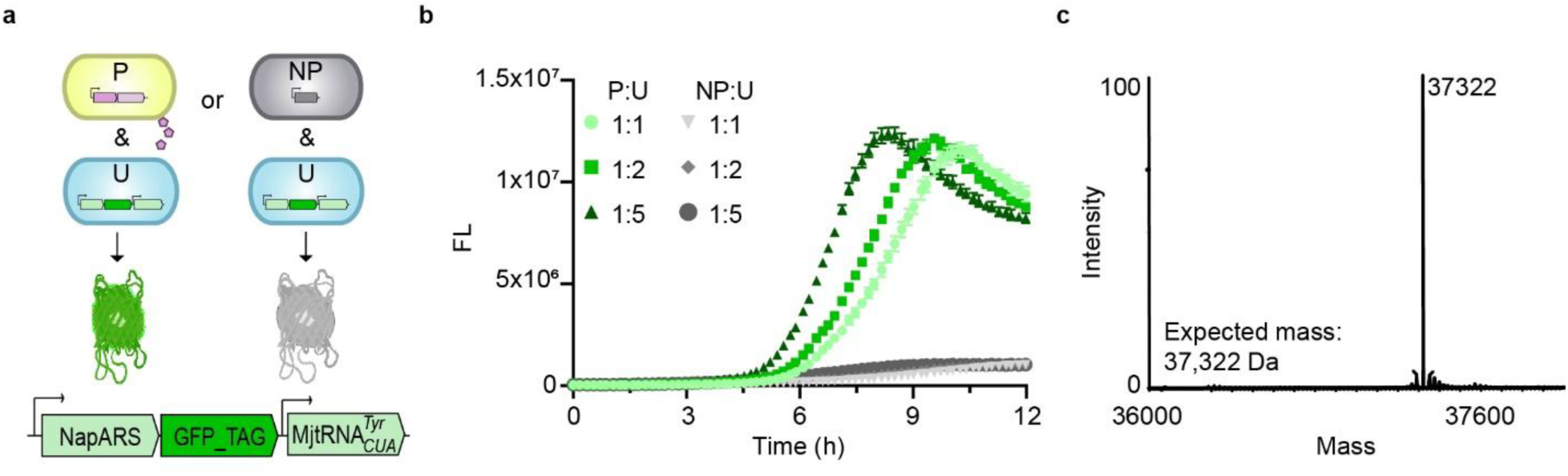
Incorporation of biosynthesized OMeTyr during a co-culture across two distinct *E. coli* strains. **a,** Predicted schema of fluorescent protein production from co-cultures that contain either an OMeTyr producer strain (P) and an OMeTyr utilizer strain (U) or an OMeTyr non-producer strain (NP) and an OMeTyr utilizer strain. The producer strain constitutively expresses *mfnG* and *aroG\**, whereas the non-producer strain does not contain OMeTyr biosynthesis genes. The utilizer strain harbors the OTS and reporting system resulting in GFP production when supplied with its nsAA. **b,** Fluorescence from co-cultures between the RR3 (DE3) strain harboring an operon containing the OTS, including the NapARS synthetase, and GFP reporter and *E. coli* strains both with and without the ability to biosynthesize OMeTyr. Strains were inoculated at varied OD_600_ ratios in MOPS EZ Rich media lacking aromatic amino acids. **c,** Mass spectra from intact protein MS of Ub-GFP with OMeTyr incorporation from metabolic synthesis in RR3 (DE3) containing an operon with the OTS, including the NapARS synthetase, and GFP reporter. Sample sizes are n=3 using biological replicates and data shown are mean ± standard deviation except for mass spectra results which represents n=1. Fluorescent outputs collected at (ex: 485 nm, em: 525 nm).

### Synthetic auxotroph dependent on OMeTyr

We therefore planned to modify our system such that the utilizer would incorporate OMeTyr into an essential protein rather than GFP. Because existing synthetic auxotrophs exhibit low or undetectable rates of escape from biocontainment, we decided to investigate previously characterized synthetic auxotrophic markers. Specifically, we investigated the ability of the previously reported *adk.d6* single-marker synthetic auxotrophic strain that was originally designed to depend on biphenylalanine (BipA)^15^ to grow on OMeTyr. Prior work has revealed the promiscuity of the BipARS synthetase harbored in this strain^29^. However, the synthetic auxotrophic marker must also accept OMeTyr and be active for the synthetic auxotroph to survive. Interestingly, we found that the *adk.d6* synthetic auxotroph maintained the same growth profile when provided OMeTyr as when provided BipA (Figs. 3a-b).

**Figure 3.**
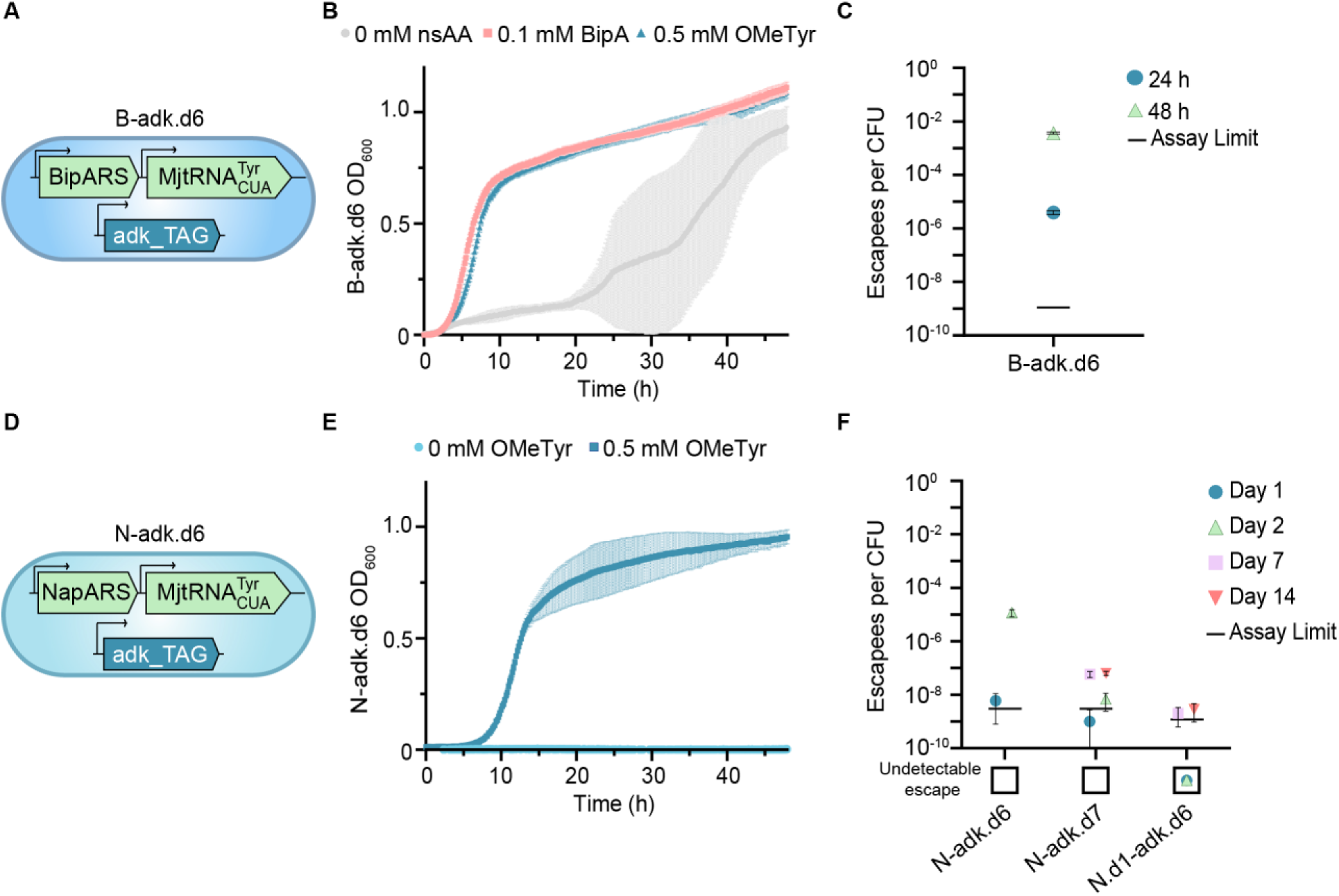
Generation and characterization of synthetic auxotrophs dependent on OMeTyr. **a,** Depiction of the previously reported B-adk.d6 synthetic auxotroph. **b,** Kinetic growth dynamics of B-adk.d6 in LB media without addition of an nsAA, with 0.1 mM BipA, or with 0.5 mM OMeTyr for 48 h at 34 °C. **c,** Escape frequency of B-adk.d6. Cells were cultured in permissive liquid media overnight, washed repeatedly, and then plated on solid media for monitoring. Colony forming units (CFUs) appearing on non-permissive plates were used to determine the number of escapees, which was divided by the total CFUs plated as determined by plating serial dilutions on permissive media. **d,** Depiction of the N-adk.d6 synthetic auxotroph. **e,** Kinetic growth dynamics of N-adk.d6 synthetic auxotroph grown in LB media with and without OMeTyr for 48 h at 34 °C. **f,** Escape frequencies of N-adk.d6, N-adk.d7, and N.d1-adk.d6 over 14 days collected by counting colonies on permissive and non-permissive solid media. N-adk.d6 escape frequency only documented through Day 2 due to an inability to distinguish colonies from increased growth. Sample sizes are n=3 using biological replicates and data shown are mean ± standard deviation.

We next sought to test the rate of escape from biocontainment for the *adk.d6* synthetic auxotroph (denoted now as “B-adk.d6”) when supplying BipA in the condition permissive for growth. We cultured cells to full density in permissive liquid media, washed cells repeatedly to remove nsAA, and then plated cells on both permissive and non-permissive (without any nsAA) solid media. The escape rate of B-adk.d6 was 4.0 x 10^-6^ escapees/CFU after 24 h of incubation on solid media (Fig. 3c). Since this level of containment is poor, we sought to improve it by replacing BipARS with NapARS. Using transformation, passaging, and selection, we replaced the plasmid harbored by B-adk.d6 with a plasmid that contained the NapARS-MjtRNA^Tyr^ OTS. These modifications generated the new N-adk.d6 synthetic auxotroph (Fig. 3d). Sequencing of N-adk.d6 revealed that its synthetase was a chimera of NapARS and BipARS (NapARS’) that exhibited comparable fold-change in FL/OD_600_ to NapARS at 24 h after inoculation (Supplementary Fig. 3).

We next investigated the ability of our new synthetic auxotroph to remain biocontained in the absence of OMeTyr by monitoring growth for 48 h in non-permissive LB media (without OMeTyr) in 96-well microplates. We were pleased to discover that, in stark contrast with the B-adk.d6 strain, the N-adk.d6 strain did not appear to escape during 48 h of culturing under these conditions (Fig. 3e). When we compared escape frequency more quantitatively in larger culture volumes, we found that N-adk.d6 resulted in 6.0 x 10^-9^ escapees/CFU after 24 h of incubation on solid media (Fig. 3f), which was reduced by three orders of magnitude compared to B-adk.d6 (Fig. 3c). However, the escape rate of N-adk.d6 after 48 h of solid media incubation was 1.2 x 10^-5^ escapees/CFU. We next used whole genome sequencing to investigate the mechanism of escape for five randomly selected escapee colonies. The results showed that one escapee had a mutational reversion of the UAG codon to UUG (leucine) in the *adk*.d6 gene, two escapees had single mutations in the *mutS* gene, and two escapees had no mutations (Supplementary Table 6).

Since sequencing results revealed that in some cases mutation was not required for escape from biocontainment, we hypothesized that instead low levels of mischarging the orthogonal tRNA with standard amino acids catalyzed by the NapARS’ may be responsible. Accordingly, we next investigated the influence of orthogonal aminoacyl-tRNA synthetase expression level on escape frequency. When we plated N-adk.d6 on non-permissive solid media that lacked both the inducer of the synthetase and the nsAA, we observed a 14-day escape rate of 3.0 x 10^-8^ escapees/CFU (Supplementary Fig. 4a). This 14-day escape rate without synthetase induction improved upon the 2-day escape rate with synthetase induction for N-adk.d6 by three orders of magnitude and was now a level of biocontainment that seemed sufficient to the investigate the design of an exclusive reliance, the focus of this study. Unfortunately, we found that the permissive condition required the presence of the inducer for growth of the bacterium (Supplementary Fig. 4b), so we considered two engineering approaches to lower the escape rate in the presence of the inducer.

To decrease the likelihood of translation of Adk.d6 in the absence of OMeTyr, we scanned the coding sequence for candidate sites to substitute a second in-frame UAG codon (Supplementary Fig. 5). We found that amino acid position 99, which was predicted to be highly permissive based on sequence conservation analysis, tolerated the substitution of an alanine to OMeTyr in what was originally an apparently spacious pocket with considerable distance from the ligand binding pocket. This substitution was performed using multiplex automatable genome engineering (MAGE)^30,31^ and resulted in the N-adk.d7 synthetic auxotroph, which exhibited a 14-day escape rate of 6.4 x 10^-8^ escapees/CFU under conditions where permissive and non-permissive media are identical other than inclusion of OMeTyr (Fig. 3f). Based on the escape rate that we had observed without synthetase induction, we also investigated whether we could reduce escape by introducing an in-frame stop codon in the NapARS’ to lower its expression level starting from the N-adk.d6 strain. We again scanned the coding sequence for candidate sites and identified a target location in a spacious loop region far from any ligand binding locations. We inserted a UAG at position 251 of the NapARS’, creating an OMeTyr-dependent NapARS’ (Supplementary Fig. 6). This strategy resulted in our best performing synthetic auxotroph, N.d1-adk.d6, which exhibited a 14-day escape rate of 2.8 x 10^-9^ escapees/CFU (Fig. 3f). While there is some heterogeneity in the measurement of escape rates from biocontainment in prior efforts due to variations in monitoring time or change of more than one variable between permissive and non-permissive conditions, the performance of our biocontainment system was now comparable to the top reported systems where any escape is still detected^32^. Moreover, we determined this level of biocontainment sufficient to proceed to testing our primary project objectives of designing an exclusive reliance.

To advance towards co-culture, we determined the specific growth rates of the N-adk.d6 and N.d1-adk.d6 strains as well as their minimum required OMeTyr concentration for growth. When 0.5 mM OMeTyr was supplied, the specific growth rates of N-adk.d6 and N.d1-adk.d6 were 0.74 h^-1^ and 0.51 h^-1^, respectively (Supplementary Fig. 7). For comparison, the specific growth rate of the OMeTyr producer strain was 0.89 h^-1^. To determine the minimum required OMeTyr concentration, we supplemented OMeTyr at concentrations as low as 0.03 mM. N-adk.d6 was able to grow at this low concentration (Supplementary Fig. 8a), whereas N.d1-adk.d6 required 0.1 mM OMeTyr (Supplementary Fig. 8b). These OMeTyr concentrations were generated by the OMeTyr producer in monoculture by the time cell densities reached 5 x 10^8^ CFU/mL and 3.43 x 10^9^ CFU/mL, respectively (Supplementary Fig. 9). These results suggested that either of the two utilizer strains may be able to grow simultaneously with the producer strain in co-culture.

### Obligate commensalism based on OMeTyr

Having engineered robust producer and utilizer strains, we next investigated our concept of orthogonal obligate commensalism within a microbial co-culture. We hypothesized that, in media lacking OMeTyr, N.d1-adk.d6 should rely on the OMeTyr producing strain (Fig. 4a). For this to be true, the N.d1-adk.d6 strain must not be able to circumvent its dependence on OMeTyr through cross-feeding of other nutrients from prototrophic *E. coli* cells, and it also must not be able to grow using the OMeTyr that it had previously been exposed to. To help address the latter requirement, we measured the persistence of N.d1-adk.d6, which is the length of time that the cells are able to survive in the absence of OMeTyr under conditions that do not generate viable escapees. When targeting the introduction of 8 x 10^6^ CFU at inoculation in liquid media lacking OMeTyr, we observed no persistence of the synthetic auxotroph at 24 h which allowed us to determine an inoculation ratio of 1:5 producer:utilizer for our co-culture (Supplementary Fig. 10). We then constructed two sets of co-cultures: (1) N.d1-adk.d6 (Kan^R^) and a non-producer *E. coli* strain (Cm^R^); (2) N.d1-adk.d6 and the OMeTyr producer strain (Cm^R^). We also included a monoculture of N.d1-adk.d6 as a negative control. In each case, we assessed the abundance of each strain using antibiotic selection: OMeTyr and Kan for the utilizer, and Cm for the producer. Since N-adk.d6 exhibited inferior biocontainment but superior fitness, we also performed experiments with N-adk.d6 in place of N.d1-adk.d6 using a 1:1 inoculation ratio for comparison (Supplementary Fig. 11).

**Figure 4.**
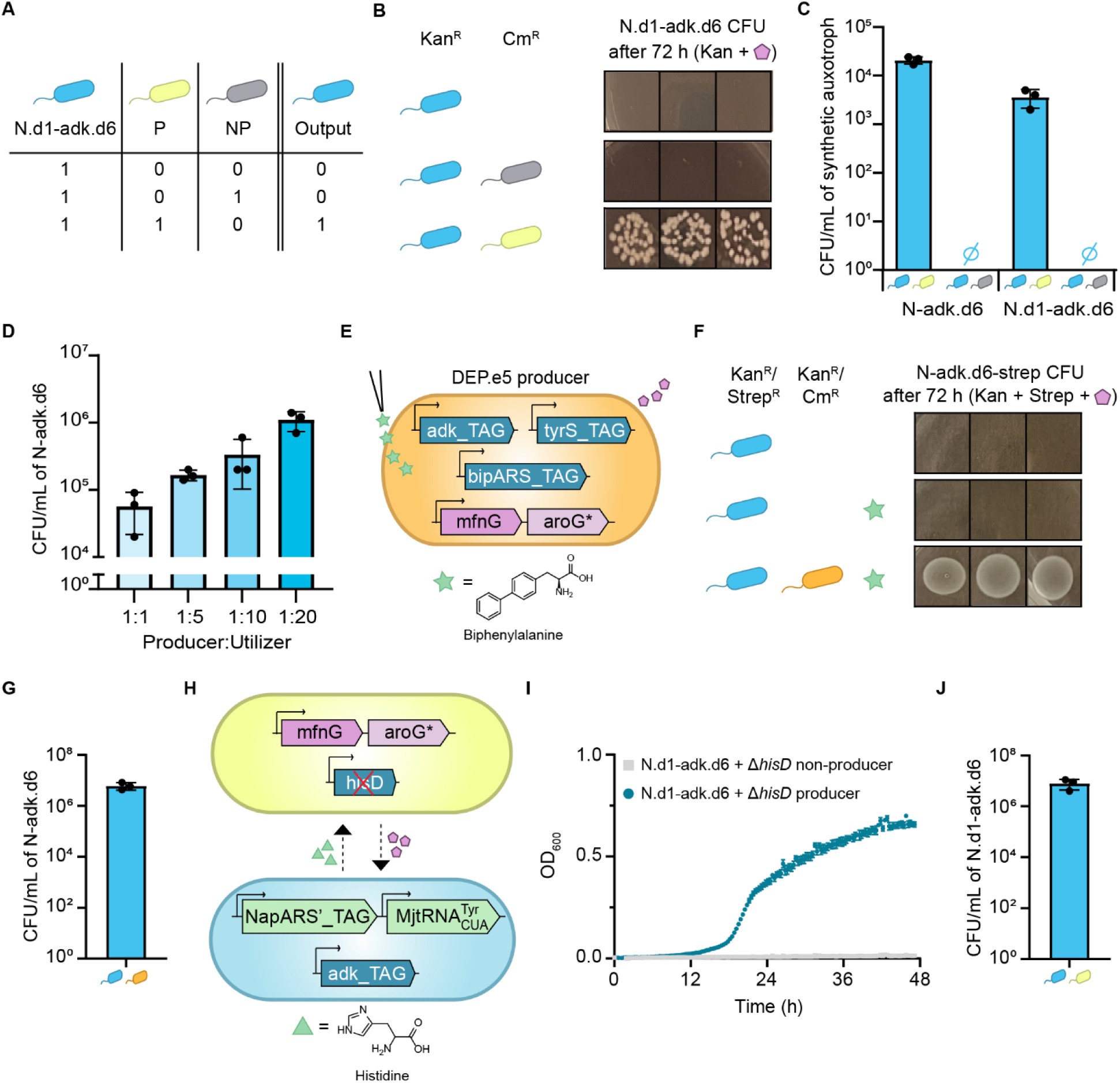
Demonstration of obligate commensalism mediated by OMeTyr. **a,** Predicted outcome of N.d1-adk.d6 survival based on various culture conditions, represented as a Boolean truth table. **b,** Co-culture results depicting n=3 replicates for each condition listed. 1^st^ row: N.d1-adk.d6 monoculture grown in non-permissive media. 2^nd^ row: N.d1-adk.d6 co-cultured with a non-producing Recoded RARE (DE3) strain of *E. coli* in non-permissive media. 3^rd^ row: N.d1-adk.d6 co-cultured with the OMeTyr producing Recoded RARE (DE3) strain of *E. coli* in non-permissive media. Images shown represent colonies 72 h after plating on solid media selective for N.d1-adk.d6. **c,** CFU/mL values of N-adk.d6 and N.d1-adk.d6 calculated from 10-fold serial dilutions of the samples grown on solid, permissive media for 96 h and 48 h, respectively following 24 h co-cultures. Values determined through colony counting from 10 µL plated media samples. **d,** Experimental CFU/mL values calculated from co-cultures of each listed producer:utilizer ratio plated on N-adk.d6 permissive media. Values determined through colony counting from 10 µL plated media samples. Sample sizes are n=3 using biological replicates and data shown are mean ± standard deviation. **e,** Depiction of the DEP synthetic auxotroph, dependent on BipA (star), with the OMeTyr production machinery. **f,** Co-culture results depicting n=3 replicates for each condition listed. 1^st^ row: N-adk.d6 monoculture in media lacking nsAAs. 2^nd^ row: N-adk.d6 monoculture in media containing 5 μM BipA externally supplied. 3^rd^ row: N-adk.d6 co-cultured with DEP capable of producing OMeTyr with 5 μM BipA externally supplied. Images shown represent colonies 72 h after plating on solid media selective for N-adk.d6. **g,** CFU/mL values of N-adk.d6 calculated from 10-fold serial dilutions of the samples grown on solid, permissive media for 24 h following a 48 h co-culture with the DEP.e5 producer in non-permissive LB liquid media. **h,** Depiction of mutualism between a histidine auxotroph, dependent on histidine (triangle), transformed to express OMeTyr biosynthetic pathway, and N.d1-adk.d6. **i,** Kinetic growth dynamics of the mutualism co-cultures with N.d1-adk.d6 and either the histidine auxotroph OMeTyr producer or the histidine auxotroph OMeTyr non-producer in M9 minimal media that lacks supplemented OMeTyr and histidine for 48 h at 34 °C. **j,** CFU/mL values of N.d1-adk.d6 calculated from 10-fold serial dilutions of the samples grown on solid, permissive media for 24 h following a 48 h co-culture with the Δ*hisD* producer in non-permissive M9 minimal media. Sample sizes are n=3 using biological replicates and data shown are mean ± standard deviation, except for images in b and f which show individual replicates.

We performed a preliminary assessment of organism-based biocontainment by plating 10 μL samples from each condition and were excited to observe that N-adk.d6 and N.d1-adk.d6 only survived in LB media after 24 h of liquid culturing with either OMeTyr supplementation as a positive control (Supplementary Fig. 11) or co-inoculation with the producer strain (Fig. 4b & Supplementary Fig. 11). We more rigorously examined the rate of escape from biocontainment for N-adk.d6 from the non-permissive liquid co-culture scenario with the non-producer by plating the entire remaining 270 μL culture volume from each triplicate well on non-permissive media (OMeTyr^-^) with antibiotic selection for N-adk.d6. We observed no CFUs throughout the entirety of the 7-day monitoring period at an assay detection limit of 7.56 x 10^-6^ escapees/CFU calculated based on the number of cells used to inoculate the co-culture (Supplementary Fig. 12). Thus, our biocontainment system functioned as desired after co-cultivation with a closely related live prototrophic *E. coli* strain, an important milestone.

To achieve our second goal of designing a system that can support utilizer strain survival in co-culture with the OMeTyr producer, we next examined co-cultures of OMeTyr producer and either the N-adk.d6 or N.d1-adk.d6 utilizer strains. We were excited to observe survival of each utilizer after 24 h of liquid culturing, with cell densities of 2.1 x 10^4^ CFU/mL and 3.7 x 10^3^ CFU/mL, respectively (Fig. 4c). Thus, we succeeded in creating the unidirectional exclusive reliance that we intended. However, for rigorous characterization about the fate of utilizer strains exposed to the producer strain, we also plated the remaining culture volume from each triplicate well on non-permissive media with antibiotic selection for the synthetic auxotroph. Here, we observed that exposure of N-adk.d6 to the producer strain facilitates considerable escape from biocontainment (Supplementary Fig. 12), whereas exposure of N.d1-adk.d6 to the producer strain only resulted in a total of 17 viable escapees across all replicates (Supplementary Fig. 13). Because this number of escapees exceeds that expected from the N.d1-adk.d6 strain alone (zero escapees), we briefly investigated whether escapees had acquired the *mfnG* containing plasmid that encoded chloramphenicol resistance from the producer strain. We found that none of them exhibited chloramphenicol resistance, which suggests that a mechanism other than horizontal gene transfer may be at play at a low but non-negligible rate. Future studies will interrogate this newly observed escape mechanism further as well as potential strategies to eliminate it.

### Control of orthogonal symbiosis culture composition

While our utilizer initially survived in the presence of producer, it did not thrive. As such, we sought to ascertain the influence of several factors on promoting co-existence rather than exclusion: Varying inoculation ratio, slowing the growth rate of the producer, and engineering mutualism. To determine the influence of utilizer growth rate and inoculation ratio on the final utilizer, we turned to the faster growing N-adk.d6 and to mathematical modeling. We modeled the observed resource competition of our co-culture system by assuming standard Monod growth kinetics for the producer strain and non-interactive, double substrate-limited growth kinetics^33^ for the utilizer (see Supplementary Methods for equations and Supplementary Tables 7-8). We estimated the model parameters using Monod growth kinetics to determine substrate-dependent growth kinetics of the producer strain, utilizer strain, and OMeTyr production (Supplementary Fig. 14). The model predicted that increasing inoculation ratio between utilizer and producer strains would yield higher utilizer abundance (Supplementary Fig. 15-18), which we experimentally verified by testing ratios of 1:1, 1:5, 1:10, and 1:20 (Fig. 4d). We were pleased to see that the use of a 1:20 inoculation ratio resulted in 1.1 x 10^6^ CFU/mL of N-adk.d6 after 24 h.

We next examined the influence of decreasing the producer growth rate by transforming a previously published synthetic auxotroph that requires BipA with the OMeTyr biosynthesis plasmid. We chose DEP.e5 as the host, which was engineered to rely on BipA and further evolved to require very low concentrations^20^. In principle, our obligate commensalism would then contain two synthetic auxotrophs that require different nsAAs, with N-adk.d6 dependent on the DEP.e5 producer for survival. We transformed the optimized plasmid for OMeTyr production into DEP.e5 to create the DEP.e5 OMeTyr producer (Cm^R^ and Kan^R^) (Fig. 4e). We found that it was able to biosynthesize above our minimum target of 0.03 mM OMeTyr within 7 h, ultimately producing 0.286 ± 0.018 mM in 24 h in its permissive LB media (supplemented with 10 µM BipA) (Supplementary Fig. 19).

Next, we investigated whether our obligate commensalism would hold between these two synthetic auxotrophs after confirming that they could not grow on the non-cognate nsAA (Supplementary Fig. 20). To continue using antibiotic selection for strain identification following a co-culture, we transformed a plasmid harboring streptomycin resistance into N-adk.d6 to generate N-adk.d6-strep (Kan^R^ and Strep^R^). Then, we prepared a co-culture of the DEP.e5 producer and N-adk.d6-strep with a 1:1 inoculation ratio in media that was permissive to DEP.e5 (5 µM BipA) but non-permissive to N-adk.d6-strep (0 µM OMeTyr). After 48 h of culturing in liquid media, we discovered via antibiotic selective plating that N-adk.d6-strep was able to survive when co-inoculated with the DEP.e5 producer (Fig. 4f). Furthermore, it could not survive in monocultures that contained BipA but lacked OMeTyr (Fig. 4f). As anticipated, when N-adk.d6-strep was co-cultured with the DEP.e5 producer, the overall abundance of N-adk.d6-strep following the co-culture increased to 6.13 x 10^6^ CFU/mL, two orders of magnitude larger than the final abundance resulting from the initial co-culture (Fig. 4g).

As another strategy to increase utilizer abundance, we sought to engineer mutualism. To construct this relationship, we used MAGE to knock out the *hisD* gene required for histidine production in *E. coli* to generate a histidine auxotroph (Δ*hisD*) and then transformed our OMeTyr production machinery into the auxotroph. Then we used a kinetic assay to monitor OD_600_ and confirm that Δ*hisD* would only grow in the presence of externally supplied histidine in M9 minimal media (Supplementary Fig. 21) and confirmed its ability to biosynthesize OMeTyr (Supplementary Fig. 22). Finally, we co-cultured both N.d1-adk.d6 and the Δ*hisD* OMeTyr producer together (Fig. 4h) in M9 minimal media and monitored OD_600_ to track growth. We were excited to see observable growth when the synthetic auxotroph and OMeTyr producing Δ*hisD* were co-cultured together and no growth when N.d1-adk.d6 was co-cultured with Δ*hisD* that lacked the OMeTyr production machinery (Fig. 4i). Following the co-culture with the Δ*hisD* OMeTyr producer, the final concentration of N.d1-adk.d6 was nearly identical to the final concentration of N.d1-adk.d6 in monoculture externally supplied with OMeTyr at 8 x 10^6^ CFU/mL and 7 x 10^6^ CFU/mL, respectively (Fig. 4j). This successful demonstration of obligate mutualism illustrates the potential for further tuning of the system to improve survival of the synthetic auxotroph when thinking about the possibility of deploying this technology.

### Orthogonal function within soil consortium

After successfully demonstrating the concept of an obligate commensalism that was orthogonal to non-producing *E. coli*, we sought to ascertain whether the system would be orthogonal to other microbes within a consortium (Fig. 5a). To begin to understand the potential for orthogonality to natural environments, we chose four microbes from a simplified maize root soil consortium^34^: *Stenotrophomonas maltophilia, Curtobacterium pusillum, Herbaspirillum frisingense,* and *Pseudomonas putida*. These four microbes exhibited growth in LB media and were susceptible to our chosen antibiotics, which allowed us to continue using antibiotic selection to monitor CFUs of the synthetic auxotroph. We first examined the orthogonality of OMeTyr production to the soil consortium with and without the OMeTyr producing *E. coli* strain present. While the microbial consortium did not produce any OMeTyr on its own, OMeTyr did appear to be consumed by the consortium since we observed that only about 0.25 mM OMeTyr remained 24 h after 0.5 mM was supplied (Fig. 5b). This biodegradability of OMeTyr is an important feature of the system if it occurs at a sufficiently slow rate because it mitigates the risk of producer-independent survival due to OMeTyr accumulation in off-target environments. When a 1:1 ratio of the OMeTyr producer:soil consortium were co-inoculated, the concentration of OMeTyr that remained in culture after 24 h was 0.05 mM (Fig. 5b). We also inoculated each of the soil microbes in pairwise co-cultures with N-adk.d6 (Kan^R^) in permissive media to screen for interactions that would exclude either microbe. We confirmed that all species were capable of co-existence after a 24 h co-culture in the N-adk.d6 permissive environment by plating on selective solid media (Supplementary Fig. 23). We noticed some light film formation from the *C. pusillum* monoculture plated with only kanamycin present, but confirmed that none of the soil microbes grew on agar containing both kanamycin and streptomycin (Supplementary Fig. 24).

**Figure 5.**
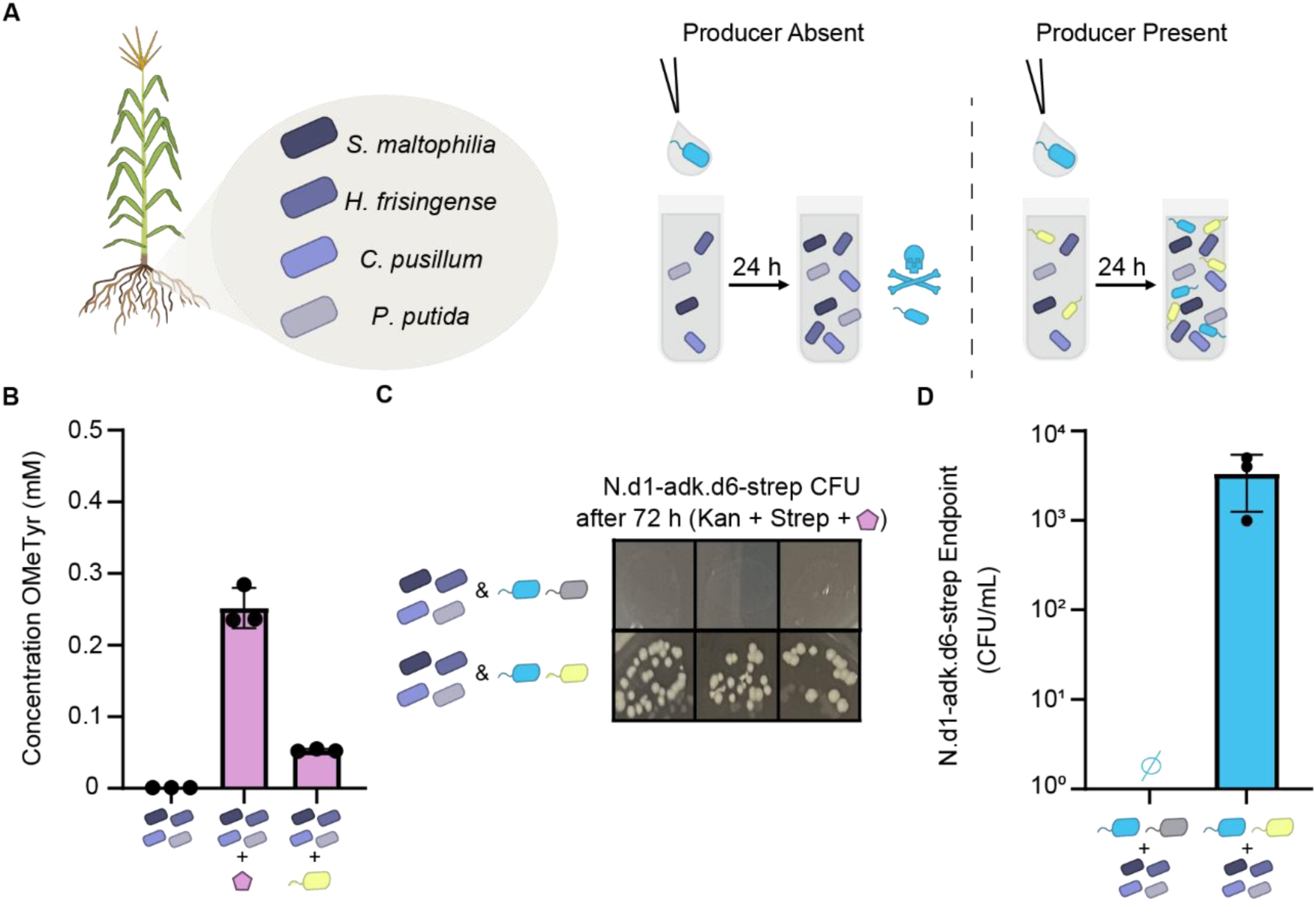
Demonstration of the orthogonality of the obligate commensalism within a microbial consortium consisting of soil isolates. **a,** Schema of desired N.d1-adk.d6 growth outcomes when grown in the absence and presence of OMeTyr producing *E. coli* when select members of a model soil consortium are also present. **b,** OMeTyr concentrations after 24 h growth of the soil consortium without externally supplied OMeTyr (left), with 0.5 mM externally supplied OMeTyr (center), and with a 1:1 inoculation ratio of the OMeTyr producing *E. coli* strain at the start of the culture (right). **c,** Growth results of N.d1-adk.d6-strep on solid, permissive media following 24 h growth in non-permissive LB in a consortium containing N.d1-adk.d6-strep, *S. maltophilia*, *C. pusillum*, *H. frisingense*, *P. putida*, and a non-producing strain of *E. coli* (top) or an OMeTyr producing strain of *E. coli* (bottom). Images shown represent colonies 72 h after plating on solid media selective for N.d1-adk.d6-strep. Sample sizes are n=3 using biological replicates each plated separately. **d,** Values of CFU/mL for N.d1-adk.d6-strep 24 h after plating cultures that were grown in non-permissive liquid media for 24 h in a consortium including the soil microbes and a non-producing (left) or OMeTyr producing (right) *E. coli* strain. Sample sizes are n=3 using biological replicates and data shown are mean ± standard deviation.

We then investigated whether N.d1-adk.d6 survival in the soil consortium would rely on inclusion of the OMeTyr producer in otherwise non-permissive media (LB). To continue using antibiotic selection for strain identification, we transformed a plasmid harboring streptomycin resistance into N.d1-adk.d6 to generate N.d1-adk.d6-strep (Kan^R^ and Strep^R^). We tested two consortium conditions: (1) Soil consortium, N.d1-adk.d6-strep, and a non-producer *E. coli* strain (1:10:1); (2) Soil consortium, N.d1-adk.d6-strep, and the OMeTyr producer strain (1:10:1). As desired, we found that N.d1-adk.d6-strep did not survive in the first consortium and did survive in the second consortium (Fig. 5c). Through dilutions we determined that there were no remaining synthetic auxotroph colonies when the OMeTyr producer was excluded from the consortium and that approximately 3.3 x 10^3^ CFU/mL N.d1-adk.d6-strep remained in the consortium that included the OMeTyr producer after 24 h (Fig. 5d). Since we did not see escape of N.d1-adk.d6 from the co-culture with the non-producer, we were interested to investigate if the presence of the soil microbes would impact its biocontainment. We examined the rate of escape of N.d1-adk.d6-strep following its addition to the consortium containing the non-producer by plating the entire remining 240 uL culture volume from each triplicate well on non-permissive media (OMeTyr^-^) with antibiotic selection for N.d1-adk.d6-strep. Again, no escape was seen during the 7-day monitoring period following the 24 h incubation in liquid culture at an assay detection limit of 2.4 x 10^-8^ escapees/CFU calculated based on the number of cells used to inoculate the consortium (Supplementary Fig. 25). Collectively, these results demonstrate that the original objectives of sustaining a biocontained microbe only in the presence of an nsAA producer were met, even in the presence of a small number of undomesticated microbes.

## Discussion

Concerns about the proliferation of genetically engineered microbes in unintended environments has limited the applications of microbial synthetic biology outside of the bioreactor. While intrinsic biological containment techniques such as synthetic auxotrophy have shown promise at permitting microbial growth in environments where an nsAA is supplied, it is unclear how to support longer-term proliferation of a contained microbe in an open environment without repeated human intervention. Here, we introduced a concept that could surmount this obstacle via autonomous establishment of a permissive environment using a second engineered organism that generates an orthogonal and essential nutrient. In doing so, we designed a form of commensalism that is both exclusive and obligate. Commensalism can be mediated by metabolic dependencies such as amino acid auxotrophy^35^ and can strongly influence the co-occurrence of diverse microbial species^36^. However, since Darwin himself questioned the existence of altruistic behavior in natural species^37^, some question whether instances of commensalism are in fact kin-selection or disguised mutualism^38,39^. Additionally, it is unknown whether there are any naturally occurring commensal extracellular species that exclusively rely on one other species. While many microbes are only culturable in the presence of other microbes^40,41^, the growth factors required to support these obligate commensals are generally not unique to one species. Some microbes initially considered exclusive obligate commensals^42–44^ appear to have been misclassified^45^. While some obligate intracellular bacterial symbionts with reduced genomes can rely exclusively on one host^46^, these interactions are less relevant to the design and understanding of dependencies between free-living organisms. Using rational design, our study reveals the molecular basis by which an exclusive obligate commensal interaction could occur in an extracellular species, an interaction not verified to occur in nature to our knowledge. Going forward, the strategies and technologies described here may aid in elucidating ecological theories while also serving as a critical linchpin for new scenarios of controlled release of engineered microbes.

Our results revealed many quantitative aspects that are critical to the success of an orthogonal obligate commensalism that features an nsAA. The rate of nsAA biosynthesis is an important variable that, in co-culture or consortium settings, must be sufficient to allow survival of the synthetic auxotroph prior to exclusion by other microbes. The required rate of nsAA biosynthesis is largely dictated by the apparent affinity of the orthogonal aminoacyl-tRNA synthetase towards a given nsAA, which in our experience can vary from as little as ten micromolar^15,47^ to several hundred micromolar concentrations^27^. Our work also demonstrated the influence of several other design parameters on promoting co-existence, such as varying inoculation ratio, controlling relative growth rates, and introducing mutualism. Successful deployment of this technology would benefit from further consideration of how to reduce metabolic burden, which relates to the parameters listed above, particularly improving affinity of the synthetase rather than increasing nsAA biosynthesis rate. Additionally, a bacterium would be less burdened if the nsAA biosynthetic genes were genome integrated, which would thereby increase the stability of this interaction.

The robust function of synthetic auxotrophy in the presence of other live microbes provided several important insights about biocontainment and our orthogonal obligate commensalism. First, we validated that our designed obligate commensalism functioned orthogonally in the presence of other living microbes, including four isolates from the wild^48^. Interestingly, we observed that the tested soil consortium consumed OMeTyr, revealing that this nsAA is readily biodegradable and is not itself orthogonal despite mediating an orthogonal commensalism. The rate of OMeTyr depletion by soil microbes is an important variable that governs the success of our microbial interaction in non-sterile environments. Additionally, the soil consortia experiments are promising given that the reliance can remain exclusive among multiple undomesticated bacteria from other genera. However, it does not guarantee orthogonality within all environmental microbiomes, and thus investigation of orthogonality in specific biomes is warranted. In particular, other untested environmental factors may interfere with the biocontainment of a synthetic auxotroph, such as bacteriophage-mediated gene transfer^49,50^. Additionally, other strategies must be pursued to limit horizontal gene transfer of engineered functions from the synthetic auxotroph, which could include introducing nsAA-dependence or more complex recoding of engineered genes, avoiding the use of plasmids, decreasing sequence homology in relevant regions, and carefully positioning antitoxin-toxin modules in the engineered host. While our strategy is designed to limit the survival of engineered microbes to intended environments, the broader ecological implications, including potential benefits or risks, should be considered.

Two eventual and illustrative deployment scenarios provide a guiding vision for current and future development of this system. One scenario is the transfer of nsAA biosynthesis functions to an engineered sterile plant that would serve as an anchor for remediation or targeted plant growth-promotion from an exclusively dependent rhizobacterium. Multiple prominent U.S. research agencies such as the Department of Energy and DARPA have invested in technology development programs towards this vision. Another scenario instead features a pair of engineered microbes with no need for plant engineering, closer to the system we report here. In this second scenario, an nsAA-producing microbe would be physically encapsulated and attached to plant seeds or a solid support adjacent to the target plant root, while the utilizer microbe would be released freely. Unlike for physically contained microbes alone, the extracapsular utilizer microbe should be capable of important behaviors such as root association and biofilm formation while surviving only in the proximity of the capsule. These envisioned applications highlight that our system offers theoretical advantages over physical containment alone (i.e., the ability of engineered microbes to roam, contact surfaces, and perform functions at those interfaces) as well as advantages over chemical-based biocontainment alone (i.e., the autonomous creation of a sustained permissive environment local to another organism). For these and other scenarios of environmental release of biocontained microbes - or for other frontier applications where species co-localization is desired, such as for synthetic endosymbionts - a vital first step is the design of an exclusive reliance of one model bacterial strain on another, which we have now demonstrated. Given the ease of OMeTyr biosynthesis through its one-step pathway from common L-tyrosine, as well as the portable nature of OTSs, this strategy should be generalizable to other, more environmentally relevant species.

## Methods

### Strains and plasmids

*Escherichia coli* strains and plasmids used are listed in **Table S1**. Molecular cloning and vector propagation were performed in DH5α. Polymerase chain reaction (PCR) based DNA replication was performed using KOD XTREME Hot Start Polymerase. Cloning was performed using Gibson Assembly with constructs and oligos for PCR amplification shown in **Table S2**. Genes were purchased as G-Blocks or gene fragments from Integrated DNA Technologies (IDT) (Coralville, IA) or Twist Bioscience (San Francisco, CA) and were optimized for *E. coli* K12 using the IDT Codon Optimization Tool with sequences shown in **Table S3** and **Table S4**. The new synthetic auxotrophs constructed in this study were derived from the published *adk.d6* and DEP.e5 synthetic auxotrophs, which were kindly provided by George Church’s laboratory at Harvard Medical School.

### Materials and Chemicals

The following compounds were purchased from MilliporeSigma (Burlington, MA, USA): kanamycin sulfate, chloramphenicol, carbenicillin disodium, streptomycin sulfate salt, potassium phosphate dibasic, potassium phosphate monobasic, M9 salts, magnesium sulfate, calcium chloride dihydrate, imidazole, glycerol, sodium dodecyl-sulfate, Tris base, HEPES, trifluoroacetic acid, L-tyrosine, L-methionine, L-tryptophan, and KOD EXTREME Hot Start polymerase. Agarose and ethanol were purchased from Alfa Aesar (Ward Hill, MA, USA). D-glucose was purchased from TCI America (Portland, OR, USA). Anhydrotetracycline (aTc) and isopropyl ß-d-1-thiogalactopyranoside (IPTG) were purchased from Cayman Chemical (Ann Arbor, MI, USA). Hydrochloric acid was purchased from RICCA (Arlington, TX, USA). Acetonitrile, sodium chloride, *O*-methyl-L-tyrosine, biphenylalanine (no longer available), LB Broth powder (Lennox), and LB Agar powder (Lennox) were purchased from Fisher Chemical (Hampton, NH, USA). L-Arabinose was purchased from VWR. A MOPS EZ-rich defined medium kit and components were purchased from Teknova (Hollister, CA, USA). Trace Elements A was purchased from Corning (Corning, NY, USA). Taq DNA ligase was purchased from GoldBio (St. Louis, MO, USA). Phusion DNA polymerase and T5 exonuclease were purchased from New England BioLabs (NEB) (Ipswich, MA, USA). Sybr Safe DNA gel stain was purchased from Invitrogen (Waltham, MA, USA). Kapa2G fast multiplex mix was purchased from Roche Diagnostics (Indianapolis, IN, USA). DNeasy Blood and Tissue Kit purchased from QIAGEN Sciences (Germantown, MD, USA).

### Biosynthesis of OMeTyr

Specified strains of *E. coli* (see Table S1) were transformed with plasmids expressing OMeTyr production pathway genes. For OMeTyr production, strains were inoculated from frozen stocks and grown overnight in 3 mL Lysogeny broth-Lennox media (LB: 10 g/L bacto tryptone, 5 g/L sodium chloride and 5 g/L yeast extract) and specified antibiotics at 34 °C with shaking at 250 RPM. These overnight cultures were used to inoculate experimental cultures targeting an OD_600_ value of 0.1. To investigate the impacts of the co-expression of aroG* on OMeTyr production, the following specified strains were inoculated in 3 mL M9 minimal media (Corning Trace Elements A (1.60 μg/mL CuSO_4_ · 5H_2_O, 863.00 μg/mL ZnSO_4_ · 7H_2_O, 17.30 μg/mL Selenite · 2Na and 1155.10 μg/mL ferric citrate), 5X M9 salts, 2 mM MgSO_4_, and 0.1 mM CaCl_2_) with 1% glucose (w/v), 0.004 mM biotin, and 0.1 μg/mL aTc. In addition, sAMF1 was grown with 1 mM IPTG, 15 μg/mL kanamycin, and 17 μg/mL chloramphenicol, sAMF2 was grown with 30 μg/mL kanamycin, and Recoded RARE (DE3) was grown with 50 μg/mL carbenicillin. Cultures were incubated at 34 °C with shaking at 250 RPM for 24 h. To investigate the impacts of a constitutive promoter on OMeTyr production, sAMF3 and sAMF4 were each inoculated in 4 mL LB with 30 μg/mL kanamycin. 0.1 μg/mL aTc was also added to the cultures containing sAMF3 at inoculation. Cultures were incubated at 34 °C with shaking at 250 RPM. Supernatant samples were taken every other hour for the first 7 h post inoculation with a final sample taken at 24 h. To compare OMeTyr production in M9 minimal media to production in LB, sAMF4 was inoculated in 4 mL M9 minimal media with 1% glucose (w/v), 0.004 mM biotin, and 30 μg/mL kanamycin and the steps for the time course described above were repeated. All biosynthesis of OMeTyr was quantified via supernatant sampling, which involved pelleting via centrifugation using an Eppendorf 5430R refrigerated centrifuge, and extracting supernatant media to be analyzed using HPLC.

### nsAA incorporation assays

For the initial synthetase screen, the orthogonal AARS/tRNA pairs, cloned with pEVOL plasmids were transformed into an *E. coli* MG1655 (DE3) strain with a pZE plasmid expressing a reporter protein fusion consisting of a ubiquitin domain, followed by an in-frame amber suppression codon, followed by GFP (pZE-Ub-AG-GFP). Sequences of the synthetases screened can be found in Table S4 and the reporter protein used in this study can be found in Table S5. These transformations were inoculated from frozen stocks and grown overnight in 3 mL LB with 30 μg/mL kanamycin and 34 μg/mL chloramphenicol at 37 °C with shaking at 250 RPM. After overnight growth, the cells were inoculated at 100x dilution at 37 °C in 300 μL MOPS EZ Rich media with aromatic amino acid (Phe, Trp, and Tyr) dropout in deep 96-well plates with a specified concentration of OMeTyr (0 mM or 1 mM), 30 μg/mL kanamycin, 34 μg/mL chloramphenicol, and 2% glucose (w/v) with shaking at 1,000 RPM and an orbital radius of 3 mm. At mid-exponential growth (OD_600_ ∼0.5), 0.1 μg/mL aTc and 0.2% (wt/vol) L-arabinose were added to induce transcription of mRNA that requires UAG suppression to form full-length GFP. Cultures were returned to 37 °C and grown for 18 h before pelleting via centrifugation. GFP fluorescence was quantified using a Spectramax i3x plate reader with Softmax Pro 7.0.3 software (excitation = 485 nm, emission = 525 nm).

For the comparison of the BipARS, NapARS, and NapARS’ aminoacyl-tRNA synthetases, strains sAMF8-10 were inoculated from frozen glycerol stocks and grown overnight in 3 mL LB media with 15 μg/mL kanamycin and 25 μg/mL carbenicillin at 34 °C with shaking at 250 RPM. Following overnight growth, the cells were used to inoculate culture tubes containing 15 μg/mL kanamycin, 25 μg/mL carbenicillin, 0.1 μg/mL aTc, 0.2% (wt/vol) L-arabinose, and either 0 or 0.5 mM OMeTyr grown again at 34 °C with shaking at 250 RPM. All three strains were inoculated in triplicate for both conditions targeting an OD_600_ of 0.05. Samples were taken 4 h, 8 h, and 24 h post inoculation, which involved removing a 300 μL sample from each culture tube and pelleting cells via centrifugation and resuspending the pellets in PBS. Both OD_600_ and GFP fluorescence were quantified using a Spectramax i3x plate reader with Softmax Pro 7.0.3 software (excitation = 485 nm, emission = 525 nm).

### HPLC analysis

OMeTyr was quantified via high-performance liquid chromatography (HPLC) using an Agilent 1100 Infinity model equipped with a Zorbax Eclipse Plus-C18 column (part number: 959701-902, 5 μm, 95Å, 2.1 x 150 mm). We used an initial mobile phase of solvent A/B = 100/0 (solvent A, 50 mM potassium phosphate, pH 7.0; solvent B, 0.1% trifluoroacetic acid in acetonitrile, flow 1 mL min^-1^) and maintained for 1 min. A gradient elution was performed (A/B) with: gradient from 100/0 to 85/15 (flow 1 mL min^-1^ to 0.5 mL min^-1^) for 1-6 min, gradient from 85/15 to 30/70 (flow 0.5 mL min^-1^) for 6-25 min, gradient from 30/70 to 100/0 (flow 0.5 mL min^-1^ to 1.0 mL min^-1^) for 25-26 min, and equilibration at 100/0 for 26-27 min. Absorption was monitored at 276 and 250 nm.

### Co-culture for GFP production

Strains sAMF4-6 were inoculated from frozen stocks and grown overnight in 3 mL MOPS EZ rich media with 30 μg/mL kanamycin and 2% glucose (w/v) at 34 °C with shaking at 250 RPM. Prior to inoculation, all overnighted cells were diluted to OD 1.0. From the diluted cultures, the strains were used to inoculate MOPS EZ rich media with aromatic amino acid (Phe, Trp, and Tyr) dropout in a 96-well plate with 30 μg/mL kanamycin, 2% glucose (w/v), 0.004 mM biotin, and 0.1 μg/mL aTc with 200 μL total working volume at indicated ratios (1:1, 1:2, and 1:5), with 2 uL of the OMeTyr (non-)producer strain used for each ratio. Cultures were grown at 34 °C using a SpectraMax i3x plate reader for 18 h, with OD_600_ and GFP fluorescence (excitation = 485 nm, emission = 525 nm) measurements taken every 10 min, with medium plate shaking between reads for 12 h.

### Purification of GFP with OMeTyr incorporation

Strains sAMF5 and sAMF7 were inoculated from frozen stocks and grown overnight in 3 mL LB containing 30 μg/mL kanamycin. The next day, these cultures were used to inoculate 150 mL of an aromatic amino acid (Phe, Trp, Tyr) dropout version of MOPS EZ rich media containing 2% glucose, 30 μg/mL kanamycin, 0.1 mM Trp, 0.004 mM biotin and 0.1 μg/mL aTc targeting an inoculation OD_600_ of 0.05 for each strain. The culture was incubated in a 34 °C shaking incubator at 250 RPM for 18 h overnight. The next day, the cells were pelleted by centrifugation using an Avanti J-15R refrigerated Beckman Coulter centrifuge at 4 °C and 4000 RPM for 10 min. Supernatant was aspirated and the pellets were resuspended in 10 mL of lysis buffer (25 mM HEPES, 10 mM imidazole, 300 mM NaCl, 10% glycerol, pH 7.4) and disrupted via sonication using a QSonica Q125 sonicator with 6 cycles of 5 s at 75% amplitude and 10 s off for 2 min cycles. The lysate was distributed into microcentrifuge tubes and centrifuged for 1 h at 18,213 *g* at 4 °C. The protein-containing supernatant was then removed and loaded into a HisTrap Ni-NTA column using an ÄKTA Pure GE FPLC system. After performing an isocratic wash at 25 mM and 50 mM imidazole, GFP was eluted in 250 mM imidazole in 1.5 mL fractions, which was clear to see via visible inspection. The fractions were combined and applied to an Amicon column (10 kDa MWCO) and the buffer was diluted at least 1,000x with a 25 mM HEPES buffer at pH 7.2. The protein was flash frozen using ultracold (−80 °C) ethanol and stored at −80 °C.

### Mass spectrometry

For intact protein MS measurements, samples were submitted to a Waters Acquity UPLC H-Class with an Acquity Protein BEH C4 column (part number: 186004495, 1.7 µm, 300Å, 2.1 x 50 mm) coupled to a Xevo G2-XS Quadrupole Time-of-Flight (QToF) mass spectrometer. Protein sample was injected into a Waters Acquity with an initial mobile phase of solvent A/B = 85/15 (solvent A, water, 0.1% formic acid; solvent B, acetonitrile, 0.1% formic acid) held at 85/15 for 1 minute followed by a gradient elution from (A/B) 85/5 to 5/95 over 5 min. Flow rate was maintained at 0.5 mL min^-1^. The QToF mass spectrometer measurements were obtained following positive electrospray ionization (ES+) with source temperature of 150 °C. The spectra were analyzed from m/z 500 to 2000 and the spectra was deconvoluted using the maximum entropy function in MassLynx software. Here, an assumed uniform gaussian distribution (width = half height of 0.60 Da with 30% intensity right and left channels) was used to evaluate a mass range from 37000 to 38000 Da (resolution: 0.1 Da/channel) at an elution time of 3.2 minutes.

For LC-MS/MS measurements, protein samples (20 µg/lane) were first submitted to SDS-PAGE. The gel was then stained with Bio-Safe^TM^ Coomassie stain (Bio-Rad), and de-stained with water. Then, protein bands corresponding to the molecular weight of the protein of interest were cut from the polyacrylamide gel and the gel fragments were digested using an in-gel tryptic digestion kit (Thermo Scientific: p/n 89871). Following digestion, extracted peptides were then desalted using Pierce^TM^ pipette tips (ThermoFisher), dried, and resuspended in 20 µL of 0.5% acetic acid (pH 4.5) and then subjected to LC-MS/MS using a Thermo Scientific Orbitrap Eclipse Tribrid Mass Spectrometer (MS; Thermo Scientific) with an Ultimate 3000 nano-liquid chromatography (nano-LC) and a FAIMS Pro Interface (Thermo Scientific). The LC-MS/MS analysis was performed using an Orbitrap Eclipse MS (Thermo Scientific). Peptides were first loaded onto a trap column (PepMap C18) and then separated by an analytical column (PepMap C18, 2.0 um; 15 cm x 75mm I.D.; Thermo Scientific) at 300 nl/min flow rate using a binary buffer system (buffer A, 0.1% formic acid in water; buffer B, 0.1% formic acid in acetonitrile) with a 165-min gradient (1% to 10% in 8 min; then to 25% buffer B over 117 min; 25% to 32% buffer B in 10 min, then to 95% buffer B over 3 min; back to 1% B in 5 min, and stay equilibration at 1% B for 20 min). Multiple CVs (−40, −55 and −75) were applied for FAIMS separation. For all experiments, the survey scans (MS1) were acquired over a mass range of 375-1500 m/z at a resolution of 60,000 in the Orbitrap. The maximum injection time was set to Dynamic, and AGC target was set to Standard. Monoisotopic peak selection was set to Peptides, and the charge state filter was set to 2-7. For MS/MS acquisition, precursors were isolated with a width of 1.6 m/z, fragmented with HCD using 30% collision energy with a maximum injection time of 100 ms, and collected in Orbitrap at a resolution of 15,000. The dynamic exclusion was set to 60 s and can be shared across different FAIMS experiments. LC-MS/MS data was collected in n=3 technical replicates.

Proteomic analysis was performed in the MaxQuant-Andromeda software suite (version 1.6.3.4) with the default parameters^51^. An *E. coli* reference proteome (strain BL21-DE3; taxonomy_id:469008) was used for database search. Other parameters include: trypsin as an enzyme with maximally two missed cleavage sites; protein N-terminal acetylation, methionine oxidation, and OMeTyr substitution for tyrosine were entered as variable modifications; cysteine carbamidomethylation as a fixed modification; and peptide length was set to a minimum of 7 amino acids. The false-discovery rate of high-confidence protein and peptide identification was 1%. Peptide intensity values derived from MaxQuant were used for quantification.

### Escape assays

For kinetic, liquid media escape assays, the strains being tested were inoculated from frozen stocks and grown overnight in 4 mL of their permissive LB media (34 μg/mL chloramphenicol, 0.005% w/v sodium dodecyl sulfate (SDS), 20 mM tris-HCl buffer, 0.2% w/v arabinose and 0.1 mM BipA for B-adk.d6 and 30 μg/mL kanamycin, 0.005% w/v sodium dodecyl sulfate (SDS), 20 mM Tris-HCl buffer, 0.1 μg/mL aTc and 0.5 mM OMeTyr for N-adk.d6) at 34°C with shaking at 250 RPM. Overnight cultures were washed by pelleting the cells through centrifugation at 4 °C, 4000 RPM for 5 min, aspirating the spent media, and resuspending the pellet in fresh LB. Following two washes, the cells were diluted to an OD_600_ of 1.0, then they were used to inoculate experimental cultures in 200 μL volumes in a 96-well plate at 100x dilution. Cell growth was monitored for 48 h in the synthetic auxotroph’s non-permissive media (permissive media lacking required nsAA) in a SpectraMax i3x plate reader with medium plate shaking at 34°C with OD_600_ readings taken every 10 min. Three biological replicates were used to visualize the escape of each strain.

For solid media escape assays, all strains were inoculated from frozen stocks and grown overnight in their specified permissive media at 34 °C with shaking at 250 RPM. Cells were then washed as described for liquid media escape assays and resuspended in fresh LB media following washes. Three biological replicates were plated on permissive media following 10-fold serial dilutions in LB to determine the number of viable CFU’s plated and monitored for at least 2 days at 34 °C. To conduct serial dilutions, cells were pipetted up and down 10 times in each new well, new pipette tips were used for each round of serial dilutions, and multiple serial dilutions for each condition were plated to ensure cells were appropriately diluted 10-fold. The cell counts from permissive plates were also used to calculate the assay limit as 1/(average total viable CFU). Three biological replicates were also plated on non-permissive media without prior serial dilutions and monitored for up to 14 days at 34 °C. Colonies were counted after the specified number of days and escape frequencies were calculated as colonies observed on non-permissive media lacking the synthetic auxotroph’s nsAA per average viable CFU plated for each strain.

### MAGE

Strain sAMF12 was inoculated from a frozen glycerol stock and grown overnight in permissive media at 34 °C with shaking at 250 RPM. The following day, a starter culture was inoculated in its permissive LB media (15 μg/mL kanamycin, 17 μg/mL chloramphenicol, 0.005% w/v sodium dodecyl sulfate (SDS), 20 mM Tris-HCl buffer, 0.1 μg/mL aTc and 0.5 mM OMeTyr) targeting an OD_600_ of 0.05 and grown at 34 °C to mid-exponential phase (OD_600_ ∼0.4). Three cycles of MAGE were performed as previously described^52^ using 5 μM of the oligonucleotide targeting adk.d6 in Table S2. Following three rounds of MAGE, colonies were screened for the introduction of the TAG codon using multiplex allele-specific colony PCR followed by sequence verification. Following sequence confirmation, passaging in permissive LB media in the absence of kanamycin was used to remove the pORTMAGE-EC1 plasmid for the creation of strain N-adk.d7. This process was repeated for strain sAMF13 using the oligonucleotide targeting *hisD* in Table S2. Following five rounds of MAGE, colonies were screened and passaged as described for the generation of strain N-adk.d7 to create the Δ*hisD* strain.

### Co-culture assays

For the co-cultures involving N-adk.d6, the strain was inoculated from its frozen stock and grown overnight in 4 mL of its permissive LB media. Strains sAMF4 and sAMF11 were also inoculated from frozen stocks and grown overnight in 4 mL LB with 34 μg/mL chloramphenicol. As described for the escape assays, prior to inoculation strains were washed and diluted to an OD_600_ of 1.0. A 96-deep-well plate with non-permissive media and no antibiotics was inoculated with 296 μL media and 4 μL culture volume. The conditions tested include a 1:1 co-culture and individual monocultures of each strain. Within the same plate we included a permissive media control for both the co-culture and N-adk.d6 monoculture. Cultures were incubated for 24 h at 34°C with shaking at 1000 RPM and an orbital radius of 3 mm. Following incubation, 10 μL samples from each well were plated on solid, permissive media containing 30 μg/mL kanamycin and solid media containing 34 μg/mL chloramphenicol to separately check for growth of N-adk.d6 or sAMF4 and sAMF11 respectively. To determine colony counts, 10-fold dilutions from the N-adk.d6 monoculture and the co-culture containing N-adk.d6 and sAMF4 were also plated on permissive media containing 30 μg/ mL kanamycin. The solid media plates were incubated at 34°C for at least 24 h prior to counting colonies. To test different ratios, the inoculation volume of the (non-)producer was always held constant at 2 μL and the inoculation volume of N-adk.d6 was increased to match the designated ratio.

For the co-culture investigation involving N.d1-adk.d6, the strain was inoculated from its frozen stock and grown overnight in 4 mL of its permissive LB media. Strains sAMF4 and sAMF11 were also inoculated from frozen stocks and grown overnight in 4 mL LB with 34 μg/mL chloramphenicol. As described for the escape assays, prior to inoculation strains were washed and diluted to an OD_600_ of 1.0. A 96-deep-well plate with non-permissive media and no antibiotics was inoculated with 295 μL media and 10 μL of N.d1-adk.d6 and 2 μL (non-)producer. The conditions tested include a 1:5 co-culture and individual monocultures of each strain. Within the same plate we included a permissive media control for both the co-culture and N-adk.d6 monoculture. Cultures were incubated for 24 h at 34 °C with shaking at 1000 RPM and an orbital radius of 3 mm. Following incubation, 10 μL samples from each well were plated on solid, permissive media containing 30 μg/mL kanamycin and solid media containing 34 μg/mL chloramphenicol to separately check for growth of N.d1-adk.d6 or sAMF4 and sAMF11, respectively. To determine colony counts, 10-fold dilutions from the N.d1-adk.d6 monoculture and the co-culture containing N.d1-adk.d6 and sAMF4 were also plated on permissive media containing 30 μg/mL kanamycin. The solid media plates were incubated at 34 °C for at least 24 h prior to counting colonies.

For the co-culture investigation between two synthetic auxotrophs, DEP.e5 was first transformed with the optimized OMeTyr production plasmid to generate the DEP.e5 producer. Then, the DEP.e5 producer and N-adk.d6-strep were inoculated from frozen stocks and grown overnight in 4 mL of their own permissive LB media (17 μg/mL chloramphenicol, 15 μg/mL kanamycin, 0.005% w/v sodium dodecyl sulfate (SDS), 20 mM tris-HCl buffer, 0.2% w/v arabinose and 0.01 mM BipA for the DEP.e5 producer and 15 μg/mL kanamycin, 47.5 μg/mL streptomycin, 0.005% w/v sodium dodecyl sulfate (SDS), 20 mM Tris-HCl buffer, 0.1 μg/mL aTc and 0.5 mM OMeTyr for N-adk.d6-strep). Prior to co-culture inoculation, cells were washed 3 times in the same manner described for the escape assays and diluted to an OD_600_ of 1.0 in fresh LB. Following washes, cells were inoculated into a 96-deep-well-plate with wells charged with 300 μL LB media without antibiotics containing 0.005% w/v sodium dodecyl sulfate (SDS), 20 mM tris-HCl buffer, 0.2% w/v arabinose, 0.1 μg/mL aTc, and 0.005 mM BipA (only when specified). A volume of 2 μL each strain was used to inoculate any well containing that strain. The culture was incubated for 48 h at 34 °C with shaking at 1000 RPM and an orbital radius of 3 mm. Following incubation, 10 μL samples from each well were plated on solid, permissive media selective for N-adk.d6 containing 15 μg/mL kanamycin, 47.5 μg/mL streptomycin. Plates were monitored at 34 °C for 72 h.

For the mutualism co-culture, N.d1-adk.d6 was inoculated from its frozen stock and grown overnight in 3 mL of its permissive LB media. Δ*hisD* and the Δ*hisD* producer strains were also inoculated from frozen stocks and grown overnight in 3 mL LB with 50 μg/mL carbenicillin and 34 μg/mL chloramphenicol respectively. As described for the escape assays, prior to inoculation strains were washed and diluted to OD_600_ of 1.0, however washes and dilutions were performed in PBS rather than LB. From the diluted cultures, the strains were used to inoculate M9 minimal media with 1% glucose (w/v), 0.004 mM biotin, 0.005% w/v sodium dodecyl sulfate (SDS), 20 mM Tris-HCl buffer, and 0.1 μg/mL aTc in a 96-well plate with 200 μL total working volume with 2 uL of each strain to create the Δ*hisD*:N.d1-adk.d6 (1:1) and the Δ*hisD* producer:N.d1-adk.d6 (1:1) co-culture conditions. Both permissive (OMeTyr^+^ or His^+^) and non-permissive (OMeTyr^-^ or His^-^) monoculture controls of the individual strains were also tested. Cultures were grown at 34°C using a SpectraMax i3x plate reader for 48 h, with OD_600_ measurements taken every 10 min, with medium plate shaking between reads. Immediately following 48 h growth on the plate reader, serial dilutions and 10 μL samples were plated on each strains’ permissive LB agar using antibiotic selection to determine colony counts.

### Consortium Experiments

Strains sAMF4, sAMF11, and N.d1-adk.d6-strep were inoculated from frozen stocks and grown overnight in 4 mL of their own permissive LB media. The soil microbes specified for this assay (*S. maltophilia, H. frisingense, C. pusillum,* and *P. putida*) were streaked on solid media from frozen stocks approximately 36 h prior to the experiment, with a single colony of each species grown overnight in 3 mL LB media at 34 °C with shaking at 250 RPM. Sub cultures of the soil microbes were grown the day of the experiment in 4 mL LB media at 34 °C with shaking at 250 RPM to at least 10^8^ cells/mL.^53^ Prior to culture inoculation, each soil microbe was diluted in fresh LB to the OD_600_ corresponding to 10^8^ cells/mL. As described for the escape assays, *E. coli* strains were washed and diluted to an OD_600_ of 1.0 and soil microbes were diluted to their OD_600_ value corresponding to 10^8^ cells/mL. The soil microbes were combined at equal volumes into a 1:1:1:1 consortium following each of their individual dilutions. A 96-deep-well-plate with N.d1-adk.d6-strep’s non-permissive LB media without antibiotics was charged with 298 μL media for monoculture controls and 278 μL media for consortium growth. The conditions tested include a 10:1:1 consortia containing 20 μL of N.d1-adk.d6-strep and 2 μL each of sAMF4, and the pre-combined soil consortium and a 10:1:1 consortium containing 20 μL of N.d1-adk.d6-strep, and 2 μL each of sAMF11, and the pre-combined soil consortium. Cultures were incubated for 24 h at 34 °C with shaking at 1000 RPM and an orbital radius of 3 mm. Following incubation, 10 μL samples from each well were plated on solid, permissive media containing both 15 μg/mL kanamycin and 47.5 μg/mL streptomycin as well as solid LB media without antibiotics to separately check for growth of N-adk.d6-strep compared to the rest of the consortium. To determine colony counts, 10-fold dilutions from the consortium triplicates were also plated on the same permissive media described above. The solid media plates were incubated at 34 °C prior to counting colonies, which were checked at 24 h and 72 h after plating. To more rigorously check for escape of the synthetic auxotroph following growth in the consortium containing the non-producer, the entire remaining culture volume from each well was washed 2x in the same way described for escape assays. Cells were resuspended in 240 μL fresh LB following each wash. After the second wash, the 240 μL culture volume was spread evenly across 3 non-permissive plates selective for N.d1-adk.d6 containing 15 μg/mL kanamycin and 47.5 μg/mL streptomycin. All plates were monitored at 34 °C for 7 days.

## Acknowledgements

We thank the Mass Spectrometry Facility at the University of Delaware for the mass spectrometry analysis that is supported by the National Institute of General Medical Sciences or the National Institutes of Health under award P20GM104316 and Y. Yu in particular for his assistance. This material is based upon work supported by The National Science Foundation Graduate Research Fellowship under Grant No. (1940700) (to A.M.F.), FFAR New Innovator Award, and the Mort Collins Foundation (to A.M.F.). We would also like to acknowledge the Kolter Lab for sharing their soil community with us as well as Amanda Rosier for providing us with these strains. We would also like to acknowledge Michelle Gee at the University of Delaware for running the MATLAB software to confirm it performs as expected and Roman Dickey for preparing an initial form of the MATLAB model.

## Contributions

A.M.K. conceived and supervised the study, and helped edit the manuscript text and figures; A.M.F designed and performed most experiments, analyzed data, prepared figures and wrote the manuscript; M.A.J. generated the model and its corresponding figures, conducted necessary experiments for parameter determination, wrote the text involving the model, and helped edit the manuscript; D.N.E. helped perform experiments for model parameter determination and synthetic auxotroph characterization, N.D.B. performed the initial MjTyrRS library screen tested in this study and helped edit the manuscript.

## Ethics declarations

### Competing interests

A.M.F., M.A.J., and A.M.K. are co-inventors on a filed provisional patent application pertaining to this work. N.D.B. and A.M.K. are co-founders of the commercial entity Nitro Biosciences.

### Declarations

Any opinion, findings, and conclusions or recommendations expressed in this material are those of the authors(s) and do not necessarily reflect the views of the National Science Foundation.

## Data Availability

The datasets generated during and/or analyzed during the current study are contained in the published article (and its Supplementary Information) and are publicly accessible via cited repositories or are available from the corresponding author upon reasonable request.

