## Supplementary Information for "Design of an exclusive obligate symbiosis for organism-based biological containment"

**References: ..... S41-S43**

#### Supplementary Methods

##### *Whole genome sequencing*

The randomly selected N-adk.d6 escapee colonies were inoculated from frozen glycerol stocks and grown overnight in non-permissive LB (30 μg/mL kanamycin, 0.005% w/v sodium dodecyl sulfate (SDS), 20 mM Tris-HCl buffer, and 0.1 μg/mL aTc) at 34 °C with shaking at 250 RPM. A DNeasy Blood and Tissue kit was used to extract and purify gDNA from each escapee. Samples were sent to SeqCenter for whole genome sequencing.

##### *Rational design for UAG insertions*

To choose the site for the second in-frame UAG insertion into adk.d6, the original amino acid sequence (with the sidechain at position 178 reverted to Leu) was analyzed via the PredictProtein[1] webserver using the SNAP2 algorithm[2] to determine potential permissible sites. In parallel, AlphaFold[3] was used to generate a PDB file of a predicted protein structure, which was compared to the initial adk gene (PDB ID: 4JZK) to confirm structural similarity and visualize where the binding pocket was likely to be. Then, candidate sites and residues within 5 Å of the proposed mutation site were qualitatively evaluated for their likelihood to accommodate a substitution to OMeTyr. The same method was used to choose the site for UAG replacement in NapARS'. When conducting the structural analysis in PyMOL, we used an overlap of the original archaeal *M. jannaschii* tyrosyl-tRNA synthetase complexed with tRNA(Tyr) and L-tyrosine (PDB ID: 1J1U).

##### *OMeTyr biosynthesis with DEP.e5 producer*

The DEP.e5 producer was inoculated from a frozen glycerol stock and grown overnight in its permissive LB (17 μg/mL chloramphenicol, 15 μg/mL kanamycin, 0.005% w/v sodium dodecyl sulfate (SDS), 20 mM Tris-HCl buffer, 0.2% w/v arabinose and 10 μM BipA) at 34 °C with shaking at 250 RPM. Prior to inoculation the next day, overnight cultures were washed by pelleting the cells via centrifugation at 4 °C, 4000 RPM for 5 min, aspirating the spent media, and resuspending the pellet in fresh LB. The washed culture was used to inoculate culture tubes containing 4 mL of its permissive LB targeting an $OD_{600}$ of 0.05. The cultures were grown at 34 °C with shaking at 250 RPM for the entirety of the experiment. Supernatant samples were taken every other hour for the first 7 h post inoculation with a final sample taken at 24 h. Approximately 300 μL was removed from the culture during each time point. Synthesis of OMeTyr was quantified via supernatant sampling, which involved pelleting via centrifugation and extracting supernatant media to be analyzed using HPLC.

##### *OMeTyr biosynthesis with ΔhisD producer*

The Δ*hisD* producer was inoculated from a frozen glycerol stock and grown overnight in LB media containing 34 μg/mL chloramphenicol at 34 °C with shaking at 250 RPM. The overnight culture was used to inoculate culture tubes containing 4 mL LB with 34 μg/mL chloramphenicol targeting an $OD_{600}$ of 0.05. The cultures were grown at 34 °C with shaking at 250 RPM for the entirety of the experiment. Supernatant samples were taken every other hour for the first 7 h post inoculation with a final sample taken at 24 h.

Approximately 300 µL was removed from the culture during each time point. Synthesis of OMeTyr was quantified via supernatant sampling, which involved pelleting via centrifugation and extracting supernatant media to be analyzed using HPLC.

*Escape assays from co-cultures*

To more rigorously test for escape of the synthetic auxotroph following the 1:1 co-culture between N-adk.d6 and (non-)producer in non-permissive LB, the entire remaining culture volume from each well was washed 2x in the same way described for escape assays. Cells were resuspended in 270 µL fresh LB following each wash. After the second wash, the 270 µL culture volume was spread evenly across 3 non-permissive plates selective for N-adk.d6 containing 30 µg/mL kanamycin. All plates were monitored at 34 °C for 7 days.

To more rigorously test for escape of the synthetic auxotroph following the 1:5 co-culture between the (non-)producer and N.d1-adk.d6 in non-permissive LB, the entire remaining culture volume from each well was washed 2x in the same way described for escape assays. Cells were resuspended in 240 µL fresh LB following each wash. After the second wash, the 240 µL culture volume was spread evenly across 3 non-permissive plates selective for N.d1-adk.d6 containing 30 µg/mL kanamycin. All plates were monitored at 34 °C for 7 days.

*Co-culture model development based on Monod growth kinetics*

The growth of the producer strain is modeled with Monod growth kinetics using an estimated $K_{s,f}$ based on the limiting nutrient source in LB and an estimated maximum growth rate for the producer based on growth kinetic studies (Supplementary Fig. 14).

$$\mu_{g,p}=\frac{\mu_{m,p}S_f}{K_{s,f}+S_f} \tag{1}$$

Because we anticipate resource competition within the co-culture that would lead to substrate depletion and cell death, we included a term for a constant death rate to determine the total growth rate, $dX_p/dt$.

$$\frac{dX_p}{dt}=X_p[\mu_{g,p}-k_{d,p}] \tag{2}$$

In addition to the growth rate of the producer strain, we used a growth-associated productivity model to represent the relationship between growth of the producer strain and the production of OMeTyr. We approximated the production rate based on the production observed in a time course to study OMeTyr production (Supplementary Fig. 1b and Supplementary Fig. 15).

$$\frac{dN}{dt}=Y_{N/X}\mu_{g,p}X_p \tag{3}$$

We further modelled the utilizer growth using non-interactive double-substrate limited growth kinetics[4], such that the model represents Monod growth kinetics but will change within the simulation depending on whether the feed source or OMeTyr is the limiting nutrient.

$$\mu_{g,u}=\begin{cases}\frac{\mu_{m,u}N}{K_N+N}, & \frac{N}{K_N}<\frac{S_f}{K_{s,f}}\\ \frac{\mu_{m,u}S_f}{K_{S,f}+S_f}, & \frac{N}{K_N}>\frac{S_f}{K_{s,f}}\end{cases} \qquad (4)$$

Once again applying a death rate for the net growth rate, we can obtain the following expression for the producer growth rate $dX_u/dt$.

$$\frac{dX_u}{dt}=X_u[\mu_{g,u}-k_{d,u}] \qquad (5)$$

Lastly, we modeled feed source consumption with a simple yield coefficient for cell mass[5] and estimated the value based on producer growth.

$$\frac{dS_f}{dt}=-\frac{1}{Y_{\frac{X}{s}}}\left(\frac{dX_p}{dt}+\frac{dX_u}{dt}\right) \qquad (6)$$

The above system of ordinary differential equations (Eqs. 1-6) was solved numerically using ode45 in MATLAB (R2022b). All parameters determined in the model are listed in Supplementary Tables 6, and 7 and Supplementary Fig. 14. The code is deposited here: https://github.com/KunjapurLab/orthogonal-co-culture.

The model was validated with a series of control conditions to ensure the expected growth behavior. The model outputs compared to collected data are in Supplementary Figure 15. Then, a series of controls for the producer:utilizer and non-producer:utilizer were performed to confirm the model behaves as expected in limiting cases as shown in Supplementary Figures 16 and 17.

#### Supplementary Tables

### Table S1. Strains and plasmids used in this study.

| Name | Relevant genotype | Source |
|---|---|---|
| *E. coli* strains | | |
| DH5α | F– Φ80*lac*ZΔM15 Δ(*lac*ZYA-argF) U169 *rec*A1 *end*A1 *hsd*R17 (rK–, mK+) *pho*A *sup*E44 λ– *thi*-1 *gyr*A96 *rel*A1 | NEB |
| C321.ΔA | C321(F1,M1,A$_0$)_ ΔprfA. ΔmutS::zeo$^R$. ΔtolC | Previous study[6] |
| MG1655 (DE3) | F- λ- *ilv*G- *rfb-50 rph-1* (λ DE3)<br>λ DE3 = λ sBamHIo ΔEcoRI-B int::(*lac*I::PlacUV5::T7 gene1) i21 Δ*nin*5 | Previous study[7] |
| Recoded RARE (DE3) | C321.ΔA<br>cyaA(L301), leuS(F613V), bamA(S763P), yeeJ(S1467P), ftsA(G124E), prfB(T246A), T49765C (folA promoter mutated to MG1655), 3815880 +C (rph reversion to MG1655)<br>ΔdkgB ΔyeaE Δ(yqhC-dkgA) ΔyahK ΔyjgB<br>DE3 integration at P21 attP integration site | Previous study[8] |
| Recoded RARE | RR (DE3)<br>*bioABCD* integration at phage 186 insertion site<br>Δexo, Δgam, Δbet, Δbla | Previous study[8] |
| sAMF1 | RR3 (DE3) harboring pZE-N$_{term}$His-MfnG and pACYC-AroG* | This study |
| sAMF2 | RR3 (DE3) harboring pZE-N$_{term}$His-MfnG | This study |
| sAMF3 | RR3 (DE3) harboring pZE- N$_{term}$His-MfnG-AroG* | This study |
| sAMF4 | RR3 (DE3) harboring pZE-P$_{Bba_J23018}$- N$_{term}$His-MfnG-AroG* | This study |
| | MG1655 (DE3) harboring pEVOL-MjTyrRS constructs #1-17 with pZE-Ub-UAG-GFP | Previous Study[9], see Table S4 |
| sAMF5 | RR3 (DE3) harboring pRepNapARS | This study |
| sAMF6 | RR3 harboring pZE-PiTTA | This study |
| sAMF7 | MG1655 (DE3) mtagBFP harboring pZE-P$_{Bba_J23018}$- N$_{term}$His-MfnG-AroG* | This study |
| B-adk.d6 | Adk.d6 synthetic auxotroph | Previous study[10] |
| N-adk.d6 | B-adk.d6 harboring pRepNapARS' without pEVOL-BipARS | This study |
| N-adk.d7 | N-adk.d6 with adk.d6 modified to include a second UAG codon at position 99 | This study |
| N.d1-adk.d6 | N-adk.d6 harboring pRepNapARS' where NapARS' is modified to include a UAG codon at position 251 | This study |
| sAMF8 | RR new harboring pRepNapARS without Ub-UAG-sfGFP and pZE-Ub-UAG-GFP-carb | This study |
| sAMF9 | RR new harboring pRepNapARS' and pZE-Ub-UAG-GFP-carb | This study |
| sAMF10 | RR new harboring pRepBipARS and pZE-Ub-UAG-GFP-carb | This study |
| sAMF11 | RR3 (DE3) harboring pRepTetRS-C11-noUAG | This study |
| N-adk.d6-strep | N-adk.d6 harboring strep res | This study |
| sAMF12 | N-adk.d6 harboring pORTMAGE-EC1 | This study |
| sAMF13 | C321.ΔA harboring pORTMAGE-EC1 | This study |
| DEP.e5 | Adk.d6, Tyrs.d8, BipARS synthetic auxotroph evolved from DEP[10] progenitor | Previous study[11] |
| DEP.e5 producer | DEP.e5 harboring pZE-N$_{term}$His-MfnG-AroG* | This study |
| Δ*hisD* | C321.ΔA ΔhisD | This study |
| Δ*hisD* producer | C321.ΔA ΔhisD harboring pZE-P$_{Bba_J23018}$- N$_{term}$His-MfnG-AroG* | This study |
| N.d1-adk.d6-strep | N.d1-adk.d6 harboring strep resistance | This study |
| Plasmids | | |
| pZE-N$_{term}$His-MfnG | ColE1 ori, Kan$^R$, Tet promoter with a codon optimized *MfnG* gene containing an N-terminal His tag | This study |
| pACYC-AroG* | pACYC-Duet-1 plasmid harboring *aro*G* gene from *E. coli* under an IPTG-inducible T7 promoter | Previous study[9] |
| pZE-N$_{term}$His-MfnG-AroG* | ColE1 ori, Kan$^R$ or Cm$^R$, Tet promoter with a codon optimized *MfnG* gene containing an N-terminal His tag and the *aro*G* gene from *E. coli* | This study |
| pZE-P$_{Bba_J23018}$- N$_{term}$His-MfnG-AroG* | ColE1 ori, Kan$^R$ or Cm$^R$, Bba_J23108 promoter with a codon optimized *MfnG* gene containing an N-terminal His tag and the *aro*G* gene from *E. coli* | This study |
| pZE-Ub-UAG-GFP | ColE1 Ori, Kan$^R$, TetR, Tet promoter with Ub-UAG-sfGFP expression cassette | Previous study[12] |
| pEVOL-MjTyrRS (1-17) | p15A Ori, Cm$^R$, AraC, AraBAD promoter harboring a modified Methanococcus jannaschii tyrosyl tRNA synthetase, and a constitutively expressed (proK promoter) amber suppressor M. jannaschii tyrosyl tRNA | See Table S4 |
| pRepNapARS | p15A Ori, Kan$^R$ or Cm$^R$, Tet promoter with Ub-UAG-sfGFP in an operon with NapARS, and a constitutively expressed (proK promoter) amber suppressor *M. jannaschii* tyrosyl tRNA. | This study, see Table S4 |
| pRepNapARS' | p15A Ori, Kan$^R$ or Cm$^R$, Tet promoter with NapARS', and a constitutively expressed (proK promoter) amber suppressor *M. jannaschii* tyrosyl tRNA. | This study, see Table S4 |
| pRepBipARS | p15A Ori, Kan$^R$ or Cm$^R$, Tet promoter with BipARS, and a constitutively expressed (proK promoter) amber suppressor *M. jannaschii* tyrosyl tRNA. | This study, see Table S4 |
| pZE-PiTTA | ColE1 ori, Kan$^R$, TetR, Tet promoter with a codon optimized piTTA gene bearing an N-terminal hexahistidine tag. | Previous study[13] |
| pRepTetRS-C11-noUAG | p15A Ori, Cm$^R$, Tet promoter with Ub-sfGFP in an operon with TetRS-C11, and a constitutively expressed (proK promoter) amber suppressor *M. jannaschii* tyrosyl tRNA. | This study |
| pCDF-Vanp-mCherry-Ub-UAG-GFP | CloDF13 ori, Strep$^R$, pVanCC promoter with mCherry-Ub-UAG-sfGFP | This study |
| pORTMAGE-EC1 | RSF1010 ori, Kan$^R$, harboring CspRecT and *mutL* (E32K) genes | Previous study[14] |

#### Table S2. Oligos used in this study

| Oligo Name | Sequence |
|---|---|
| MfnG ins fwd | gccatcaccatcatcaccacATGACCCCTGAGGGTAAC |
| N term His rev | gtggtgatgatggtgatgg |
| MfnG seq | GGGGTACAGGTTATGATCTT |
| MfnG ins rev | ccatgggatcccccatcaagTTACTGACCCTCCGGC |
| pZE bb fwd | cttgatgggggatcccatg |
| AroG* ins fwd | cggccgctaataaaaggagatataccatgaattatcagaacgacgatttac |
| AroG* ins rev | ccatgggatcccccatcaagttacccgcgacgcg |
| MfnG ins rev2 | ggtatatctccttttattagcggccgTTACTGACCCTCCGGCAC |
| MfnG linker AroG* seq | GAATGGTTCACAGCGGC |
| Kan res fwd | gctcttcgtccagatcatcctgatcgacaagaccggc |
| Kan res rev | gaagccggtcttgtcgatcaggatgatctggacgaagag |
| Bba_J23018 fwd | ctgacagctagctcagtcctaggtataatgctagcgaattcattaaagaggagaaaggt |
| Bba_J23108 rev | gctagcattatacctaggactgagctagctgtcagctcgaggtgaagacgaaagg |
| pRepSynth (no GFP) fwd | ttaaagaggagaaaggtaccaagcttgatgggggatcc |
| pRepSynth (no GFP) rev | tgggatcccccatcaagcttggtacctttctcctctttaatgaa |
| pEVOL seq | gaagtcagccccatacg |
| CM res fwd | caataactgccttaaaaaaattacgccccgccctg |
| pZE no res rev | tttttttaaggcagttattggtg |
| CM res rev | caggagctaaggaagctaaaatggagaaaaaaatcactggatatacc |
| pZE no res fwd | tttagcttccttagctcctg |
| UAG adk.d6A99 | C*G*TCCGGTACGTCGAATTCCAGAACGTAATCAgCATTGATGCCCtaTTCTTTCATCGCGTCTGCCTGCGGAATGGTACGCGGGAAGCCGT |
| UAG hisD KO | T*G*AGCTTTAACACAATCATTGACTGGAATAGCTGTACTGCGTAGTAATGACGCCAGCTGTTAATGCGCCCGGCGATTTCCGCCTCTGAAA |
| UAG napARS'Y251 FWD | cgaaatacttcctggaaTAGccgctgaccatcaaacg |
| UAG napARS'Y251 REV | cagcggCTAttccaggaagtatttcgcg |
| HisD seq fwd | TCGCCCTGCTGCCAG |
| HisD MASC rev | CGGCAGTTTTTGCGCAGTG |
| HisD MASC wt fwd | attgactggaatagctgtactgcgg |
| HisD MASC mut fwd | attgactggaatagctgtactgcgt |
| Adk99 MASC rev | CTTTGGAaacGTAGCCGATCTA |
| Adk99 MASC wt fwd | AGACGCGATGAAAGAAGCG |
| Adk99 MASC mut fwd | GCAGACGCGATGAAAGAATAG |

* = Phosphorothioate bonds between bases

#### Table S3. DNA G-Block used for MfnG cloning in this study.

| Oligo Name | Protein Accession Number | Sequence |
|---|---|---|
| **MfnG** | AJV88379 | ATGACCCCTGAGGGTAACGTGTCGCTTGTTGATGAGTCACTGTTGGTAGGTGTGACTGATGAGGATCGTGCGGTGCGT<br>AGCGCTCACCAGTTTTACGAACGTTTGATTGGCTTGTGGGCTCCGGCGGTCATGGAGGCCGCTCACGAACTGGGTGTCT<br>TTGCGGCGCTGGCCGAAGCTCCGGCCGACAGCGGTGAATTGGCTCGTCGCCTGGACTGCGACGCTCGCGCAATGCGCG<br>TCTTACTGGATGCACTGTACGCCTATGATGTTATCGACCGGATTCATGATACTAACGGCTTTCGTTATCTGCTGAGTGCT<br>GAAGCGCGTGAGTGCTTACTGCCAGGTACCTTGTTCTCGTTGGTGGGCAAATTCATGCATGATATTAACGTCGCGTGGC<br>CGGCATGGCGTAACCTGGCGGAAGTAGTGCGTCACGGCGCGCGCGATACTTCAGGTGCCGAATCCCCGAATGGGATCG<br>CGCAGGAAGATTACGAATCTCTTGTGGGAGGGATTAACTTCTGGGCCCCTCCTATTGTGACCACCCTGTCGCGTAAACT<br>GCGTGCTAGTGGTCGCAGCGGTGATGCTACGGCCTCAGTTTTGGATGTTGGTTGTGGCACCGGACTGTATAGCCAATTG<br>TTACTCCGTGAATTCCCCCGTTGGACGGCTACTGGGCTCGATGTGGAGCGCATTGCGACACTCGCCAATGCGCAAGCA<br>CTGCGGCTGGGCGTCGAAGAACGGTTCGCGACGCGTGCGGGTGACTTTTGGCGTGGCGGTTGGGGTACAGGTTATGAT<br>CTTGTGCTGTTCGCGAATATTTTCCACCTGCAGACCCCAGCATCAGCAGTTCGCCTGATGCGTCACGCGGCAGCCTGTC<br>TGGCTCCAGATGGGTTGGTCGCTGTGGTTGACCAAATTGTGGATGCGGATCGTGAACCAAAAACCCCACAAGATCGCT<br>TTGCGTTGTTATTCGCAGCCTCGATGACCAATACGGGCGGCGGCGATGCATACACATTTCAGGAATATGAAGAATGGT<br>TCACAGCGGCCGGTTTACAGCGGATTGAAACTTTAGATACTCCCATGCACCGCATTCTGTTGGCTCGCCGTGCGACGGA<br>ACCGTCTGCTGTGCCGGAGGGTCAGTAA |

#### Table S4. Sequences of AARS variants used in this study

| | AARS | AA Sequence |
|---|---|---|
| MjTyrRS 1 | BrbRS3[15] | MDEFEMIKRNTSEIISEEELREVLKKDEKSAGIGFEPS<br>GKIHLGHYLQIKKMIDLQNAGFDIIIELADLHAYLNQKG<br>ELDEIRKIGDYNKKVFEAMGLKAKYVYGSEWMLDKD<br>YTLNVYRLALKTTLKRARRSMELIAREDENPKVAEVIY<br>PIMQVNDIHYLGVDVAVGGMEQRKIHMLARELLPKKV<br>VCIHNPVLTGLDGEGKMSSSKGNFIAVDDSPEEIRAKI<br>KKAYCPAGVVEGNPIMEIAKYFLEYPLTIKRPEKFGGD<br>LTVNSYEELESLFKNKELHPMDLKNAVAEELIKILEPIR<br>KRL* |
| MjTyrRS 2 | bpaRS_mut[16] | MDEFEMIKRNTSEIISEEELREVLKKDEKSAGIGFEPS<br>GKIHLGHYLQIKKMIDLQNAGFDIIILLADLHAYLNQKG<br>ELDEIRKIGDYNKKVFEAMGLKAKYVYGSPFQLDKDY<br>TLNVYRLALKTTLKRARRSMELIAREDENPKVAEVIYPI<br>MQVNTSHYLGVDVAVGGMEQRKIHMLARELLPKKVV<br>MIHNPVLTGLDGEGKMSSSKGNFIAVDDSPEEIRAKIK<br>KAYCPAGVVEGNPIMEIAKYFLEYPLTIKRPEKFGGDL<br>TVNSYEELESLFKNKELHPMDLKNAVAEELIKILEPIRK<br>RL* |
| MjTyrRS 3 | DopaRS[17] | MDEFEMIKRNTSEIISEEELREVLKKDEKSALIGFEPS<br>GKIHLGHYLQIKKMIDLQNAGFDIIILLSDLNAYLNQKG<br>ELDEIRKIGDYNKKVFEAMGLKAKYVYGSEFQLDKDY<br>TLNVYRLALKTTLKRARRSMELIAREDENPKVAEVIYPI<br>MQVNDIHYLGVDVQVGGMEQRKIHMLARELLPKKVV<br>CIHNPVLTGLDGEGKMSSSKGNFIAVDDSPEEIRAKIK<br>KAYCPAGVVEGNPIMEIAKYFLEYPLTIKRPEKFGGDL<br>TVNSYEELESLFKNKELHPMDLKNAVAEELIKILEPIRK<br>RL* |
| MjTyrRS 4 | BrbRS1[15] | MDEFEMIKRNTSEIISEEELREVLKKDEKSAGIGFEPS<br>GKIHLGHYLQIKKMIDLQNAGFDIIIELADLHAYLNQKG<br>ELDEIRKIGDYNKKVFEAMGLKAKYVYGSEFQLDKDY<br>TLNVYRLALKTTLKRARRSMELIAREDENPKVAEVIYPI<br>MQVNGCHYLGVDVAVGGMEQRKIHMLARELLPKKVV<br>CIHNPVLTGLDGEGKMSSSKGNFIAVDDSPEEIRAKIK<br>KAYCPAGVVEGNPIMEIAKYFLEYPLTIKRPEKFGGDL<br>TVNSYEELESLFKNKELHPMDLKNAVAEELIKILEPIRK<br>RL* |
| MjTyrRS 5 | pmmfFRS_A65V/ S158A[18] | MDEFEMIKRNTSEIISEEELREVLKKDEKSAVIGFEPS<br>GKIHLGHYLQIKKMIDLQNAGFDIIIVLADLHAYLNQKG<br>ELDEIRKIGDYNKKVFEAMGLKAKYVYGSEWMLDKD<br>YTLNVYRLALKTTLKRARRSMELIAREDENPKVAEVIY<br>PIMQVNAMHYLGVDVAVGGMEQRKIHMLARELLPKK<br>VVCIHNPVLTGLDGEGKMSSSKGNFIAVDDSPEEIRA<br>KIKKAYCPAGVVEGNPIMEIAKYFLEYPLTIKRPEKFG<br>GDLTVNSYEELESLFKNKELHPMDLKNAVAEELIKILE<br>PIRKRL* |
| MjTyrRS 6 | Bibaf-G2[19] | MDEFEMIKRNTSEIISEEELREVLKKDEKSAGIGFEPS<br>GKIHLGHYLQIKKMIDLQNAGFDIIIELADLHAYLNQKG<br>ELDEIRKIGDYNKKVFEAMGLKAKYVYGSEWMLDKD<br>YTLNVYRLALKTTLKRARRSMELIAREDENPKVAEVIY<br>PIMQVNSIHYKGVDVAVGGMEQRKIHMLARELLPKKV<br>VCIHNPVLTGLDGEGKMSSSKGNFIAVDDSPEEIRAKI<br>KKAYCPAGVVEGNPIMEIAKYFLEYPLTIKRPEKFGGD<br>LTVNSYEELESLFKNKELHPMDLKNAVAEELIKILEPIR<br>KRL* |
| MjTyrRS 7 | pAcFRS.2.t1[20] | MDEFEMIKRNTSEIISEEELREVLKKDEKSALIGFEPS<br>GKIHLGHYLQIKKMIDLQNAGFDIIIVLADLHAYLNQKG<br>ELDEIRKIGDYNKKVFEAMGLKAKYVYGSEFQLDKDY<br>TLNVYRLALKTTLKRARRSMELIAREDENPKVAEVIYPI<br>MQVNGCHYRGVDVDVGGMEQRKIHMLARELLPKKV<br>VCIHNPVLTGLDGEGKMSSSKGNFIAVDDSPEEIRAKI<br>KKAYCPAGVVEGNPIMEIAKYFLEYPLTIKGPEKFGGD<br>LTVNSYEELESLFKNKELHPMRLKNAVAEELIKILEPIR<br>KRL* |

| | | |
|---|---|---|
| MjTyrRS 8 | pAzFRS[21] | MDEFEMIKRNTSEIISEEELREVLKKDEKSALIGFEPS GKIHLGHYLQIKKMIDLQNAGFDIIILLADLHAYLNQKG ELDEIRKIGDYNKKVFEAMGLKAKYVYGSEFQLDKDY TLNVYRLALKTTLKRARRSMELIAREDENPKVAEVIYPI MQVNVMHYDGVDVYVGGMEQRKIHMLARELLPKKV VCIHNPVLTGLDGEGKMSSSKGNFIAVDDSPEEIRAKI KKAYCPAGVVEGNPIMEIAKYFLEYPLTIKGPEKFGGD LTVNSYEELESLFKNKELHPMRLKNAVAEELIKILEPIR KRL* |
| MjTyrRS 9 | BipARS[22] | MDEFEMIKRNTSEIISEEELREVLKKDEKSAHIGFEPS GKIHLGHYLQIKKMIDLQNAGFDIIIHLADLHAYLNQKG ELDEIRKIGDYNKKVFEAMGLKAKYVYGSEWMLDKD YTLNVYRLALKTTLKRARRSMELIAREDENPKVAEVIY PIMQVNGIHYKGVDVAVGGMEQRKIHMLARELLPKKV VCIHNPVLTGLDGEGKMSSSKGNFIAVDDSPEEIRAKI KKAYCPAGVVEGNPIMEIAKYFLEYPLTIKRPEKFGGD LTVNSYEELESLFKNKELHPMDLKNAVAEELIKILEPIR KRL* |
| MjTyrRS 10 | FnYRS[23] | MDEFEMIKRNTSEIISEEELREVLKKDEKSALIGFEPS GKIHLGHYLQIKKMIDLQNAGFDIIIGLADLNAYLNQKG ELDEIRKIGDYNKKVFEAMGLKAKYVYGSEFQLDKDY TLNVYRLALKTTLKRARRSMELIAREDENPKVAEVIYPI MQVNSYHYHGVDVAVGGMEQRKIHMLARELLPKKVV CIHNPVLTGLDGEGKMSSSKGNFIAVDDSPEEIRAKIK KAYCPAGVVEGNPIMEIAKYFLEYPLTIKRPEKFGGDL TVNSYEELESLFKNKELHPMDLKNAVAEELIKILEPIRK RL* |
| MjTyrRS 11 | pCNFRS[24] | MDEFEMIKRNTSEIISEEELREVLKKDEKSALIGFEPS GKIHLGHYLQIKKMIDLQNAGFDIIIVLADLHAYLNQKG ELDEIRKIGDYNKKVFEAMGLKAKYVYGSEWMLDKD YTLNVYRLALKTTLKRARRSMELIAREDENPKVAEVIY PIMQVNGAHYLGVDVAVGGMEQRKIHMLARELLPKK VVCIHNPVLTGLDGEGKMSSSKGNFIAVDDSPEEIRA KIKKAYCPAGVVEGNPIMEIAKYFLEYPLTIKRPEKFG GDLTVNSYEELESLFKNKELHPMDLKNAVAEELIKILE PIRKRL* |
| MjTyrRS 12 | CouRS-D8[25] | MDEFEMIKRNTSEIISEEELREVLKKDEKSAEIGFEPS GKIHLGHYLQIKKMIDLQNAGFDIIIHLGDLGAYLNQKG ELDEIRKIGDYNKKVFEAMGLKAKYVYGSEYHLDKDY TLNVYRLALKTTLKRARRSMELIAREDENPKVAEVIYPI MQVNGIHYGGVDVAVGGMEQRKIHMLARELLPKKVV CIHNPVLTGLDGEGKMSSSKGNFIAVDDSPEEIRAKIK KAYCPAGVVEGNPIMEIAKYFLEYPLTIKRPEKFGGDL TVNSYEELESLFKNKELHPMDLKNAVAEELIKILEPIRK RL* |
| MjTyrRS 13 | pydtRSa2[26] | MDEFEMIKRNTSEIISEEELREVLKKDEKSALIGFEPS GKIHLGHYLQIKKMIDLQNAGFDIIIVLADLAAYLNQKG ELDEIRKIGDYNKKVFEAMGLKAKYVYGSEWSLDKDY TLNVYRLALKTTLKRARRSMELIAREDENPKVAEVIYPI MQVNVAHYVGVDVAVGGMEQRKIHMLARELLPKKVV CIHNPVLTGLDGEGKMSSSKGNFIAVDDSPEEIRAKIK KAYCPAGVVEGNPIMEIAKYFLEYPLTIKRPEKFGGDL TVNSYEELESLFKNKELHPMDLKNAVAEELIKILEPIRK RL* |
| MjTyrRS 14 | NapARS[27] | MDEFEMIKRNTSEIISEEELREVLKKDEKSALIGFEPS GKIHLGHYLQIKKMIDLQNAGFDIIILLADLHAYLNQKG ELDEIRKIGDYNKKVFEAMGLKAKYVYGSEFQLDKDY TLNVYRLALKTTLKRARRSMELIAREDENPKVAEVIYPI MQVNPAHYQGVDVVVGGMEQRKIHMLARELLPKKV VCIHNPVLTGLDGEGKMSSSKGNFIAVDDSPEEIRAKI KKAYCPAGVVEGNPIMEIAKYFLEYPLTIKRPEKFGGD LTVNSYEELESLFKNKELHPMRLKNAVAEELIKILEPIR KRL* |
| MjTyrRS 15 | pAcFRS[28] | MDEFEMIKRNTSEIISEEELREVLKKDEKSALIGFEPS GKIHLGHYLQIKKMIDLQNAGFDIIILLADLHAYLNQKG ELDEIRKIGDYNKKVFEAMGLKAKYVYGSEFQLDKDY TLNVYRLALKTTLKRARRSMELIAREDENPKVAEVIYPI MQVNGCHYRGVDVAVGGMEQRKIHMLARELLPKKV VCIHNPVLTGLDGEGKMSSSKGNFIAVDDSPEEIRAKI KKAYCPAGVVEGNPIMEIAKYFLEYPLTIKRPEKFGGD LTVNSYEELESLFKNKELHPMDLKNAVAEELIKILEPIR KRL* |

| | | |
|---|---|---|
| MjTyrRS 16 | pNFRS[29] | MDEFEMIKRNTSEIISEEELREVLKKDEKSALIGFEPS<br>GKIHLGHYLQIKKMIDLQNAGFDIIILLADLHAYLNQKG<br>ELDEIRKIGDYNKKVFEAMGLKAKYVYGSSFQLDKDY<br>TLNVYRLALKTTLKRARRSMELIAREDENPKVAEVIYPI<br>MQVNPLNYEGVDVAVGGMEQRKIHMLARELLPKKVV<br>CIHNPVLTGLDGEGKMSSSKGNFIAVDDSPEEIRAKIK<br>KAYCPAGVVEGNPIMEIAKYFLEYPLTIKRPEKFGGDL<br>TVNSYEELESLFKNKELHPMDLKNAVAEELIKILEPIRK<br>RL* |
| MjTyrRS 17 | pAFRS[30] | MDEFEMIKRNTSEIISEEELREVLKKDEKSATIGFEPS<br>GKIHLGHYLQIKKMIDLQNAGFDIIILLADLHAYLNQKG<br>ELDEIRKIGDYNKKVFEAMGLKAKYVYGSTFQLDKDY<br>TLNVYRLALKTTLKRARRSMELIAREDENPKVAEVIYPI<br>MQVNPLHYAGVDVAVGGMEQRKIHMLARELLPKKVV<br>CIHNPVLTGLDGEGKMSSSKGNFIAVDDSPEEIRAKIK<br>KAYCPAGVVEGNPIMEIAKYFLEYPLTIKRPEKFGGDL<br>TVNSYEELESLFKNKELHPMDLKNAVAEELIKILEPIRK<br>RL* |
| MjTyrRS 18 | tetRS-C11[31] | MDEFEMIKRNTSEIISEEELREVLKKDEKSAAIGFEPS<br>GKIHLGHYLQIKKMIDLQNAGFDIIIVLADLHAYLNQKG<br>ELDEIRKIGDYNKKVFEAMGLKAKYVYGSEDHLDKDY<br>TLNVYRLALKTTLKRARRSMELIAREDENPKVAEVIYPI<br>MQVNGIHYSGVDVAVGGMEQRKIHMLARELLPKKVV<br>CIHNPVLTGLDGEGKMSSSKGNFIAVDDSPEEIRAKIK<br>KAYCPAGVVEGNPIMEIAKYFLEYPLTIKRPEKFGGDL<br>TVNSYEELESLFKNKELHPMDLKNAVAEELIKILEPIRK<br>RL* |
| MjTyrRS 19 | NapARS' | MDEFEMIKRNTSEIISEEELREVLKKDEKSALIGFEPSGKI<br>HLGHYLQIKKMIDLQNAGFDIIILLADLHAYLNQKGELD<br>EIRKIGDYNKKVFEAMGLKAKYVYGSEWMLDKDYTLN<br>VYRLALKTTLKRARRSMELIAREDENPKVAEVIYPIMQ<br>VNGIHYKGVDVAVGGMEQRKIHMLARELLPKKVVCIH<br>NPVLTGLDGEGKMSSSKGNFIAVDDSPEEIRAKIKKAYC<br>PAGVVEGNPIMEIAKYFLEYPLTIKRPEKFGGDLTVNSY<br>EELESLFKNKELHPMRLKNAVAEELIKILEPIRKRL* |

#### Table S5. Sequences of reporter protein used in this study.

| Reporter Plasmid | DNA CDS | Amino Acid Sequence (*: TAG codon) |
|---|---|---|
| pZE-Ub-UAG-GFP<br><br>and<br><br>pRepAARS<br><br>with Ub-UAG-GFP reporters | ATGCAGATTTTTGTGAAGACTTTAACAGGTAAGACG<br>ATTACCCTGGAGGTGGAGTCCTCGGACACCATCGAT<br>AATGTAAAATCAAAAATCCAAGATAAGGAAGGAAT<br>CCCTCCAGACCAGCAACGTCTGATTTTCGCAGGTAA<br>ACAACTGGAGGATGGTCGCACGCTTTCGGACTACAA<br>CATCCAGAAAGAATCTACCCTTCATTTGGTTCTGCG<br>TCTGCGTGGAGGATAGTTGTTTGTGCAGGAGCTTGC<br>ATCCAAGGGCGAGGAGCTCTTTACTGGCGTAGTACC<br>AATTCTCGTAGAGCTCGATGGCGATGTAAATGGCCA<br>TAAGTTTTCCGTACGCGGCGAGGGCGAGGGCGATGC<br>AACTAACGGCAAGCTCACTCTCAAGTTTATTTGTAC<br>TACTGGCAAGCTCCCAGTACCATGGCCAACTCTCGT<br>AACTACTCTGACCTATGGCGTACAATGTTTTTCCCGC<br>TATCCAGATCACATGAAGCAACATGATTTTTTTAAG<br>TCCGCAATGCCAGAGGGCTATGTACAAGAGCGCACT<br>ATTAGCTTTAAGGATGATGGCACCTATAAGACTCGC<br>GCAGAGGTAAAGTTTGAGGGCGATACTCTCGTAAAT<br>CGCATTGAGCTCAAGGGCATTGAT**TTT**AAGGAGGAT<br>GGCAATATTCTCGGCCATAAGCTGGAGTATAATTTC<br>AATTCCCATAATGTATACATTACCGCAGATAAGCAA<br>AAGAATGGCATTAAGGCGAATTTTAAGATTCGCCAT<br>AATGTGGAGGATGGCTCCGTACAACTCGCAGATCAT<br>TATCAACAAAATACTCCAATTGGCGATGGCCCAGTA<br>CTCCTCCCAGATAATCATTATCTCTCCACTCAATCCG<br>TGCTCTCCAAAGATCCAAATGAGAAGCGCGATCACA<br>TGGTACTCCTGGAGTTTGTAACTGCAGCAGGCATTA<br>CTCATGGCATGGATGAGCTCTATAAGCTCGAGCACC<br>ACCACCACCACCACCACTAA | MQIFVKTLTGKTITLEVESSDTIDNVKSKIQDKEGIPPDQQR<br>LIFAGKQLEDGRTLSDYNIQKESTLHLVLRLRGG*LFVQEL<br>ASKGEELFTGVVPILVELDGDVNGHKFSVRGEGEGDATNG<br>KLTLKFICTTGKLPVPWPTLVTTLTYGVQCFSRYPDHMKQ<br>HDFFKSAMPEGYVQERTISFKDDGTYKTRAEVKFEGDTLV<br>NRIELKGID**F**KEDGNILGHKLEYNFNSHNVYITADKQKNGI<br>KANFKIRHNVEDGSVQLADHYQQNTPIGDGPVLLPDNHYL<br>STQSVLSKDPNEKRDHMVLLEFVTAAGITHGMDELYKLEH<br>HHHHHH |

**Table S6. Escape mechanisms for N-adk.d6 via whole-genome sequencing**

| Strain name | Media condition | Escape mechanism |
|---|---|---|
| N-adk.d6_esc-1 | Non-permissive | UAG reversion to UUG |
| N-adk.d6_esc-2 | Non-permissive | mutS (+AG @ nt 820) |
| N-adk.d6_esc-3 | Non-permissive | n/a |
| N-adk.d6_esc-4 | Non-permissive | n/a |
| N-adk.d6_esc-5 | Non-permissive | mutS (+AG @ nt 820) |

**Table S7. Parameters for Orthogonal Obligate Commensalism Co-culture Model**

| Parameters | Value | Units | Source | Purpose |
|---|---|---|---|---|
| $\mu_{m,u}$ | 0.88 | $hr^{-1}$ | Calculated (SI Fig. 10) | Max growth rate: utilizer |
| $\mu_{m,p}$ | 0.95 | $hr^{-1}$ | Estimated | Max growth rate: producer |
| $K_N$ | 19.5 | mg/L | Calculated (SI Fig. 10) | "half-rate constant" for OMeTyr |
| $K_{S,f}$ | 0.5 | mg/L | Estimated | "half-rate constant" for feed substrate |
| $k_{d,u}$ | 0.2 | hr | Estimated | Death rate for the utilizer |
| $k_{d,p}$ | 0.15 | hr | Estimated | Death rate for the producer |
| $Y_{N/X}$ | 0.125 | mg/mg | Estimated (SI Fig. 1c) | OMeTyr production yield coefficient |
| $Y_{X/S}$ | 0.4 | mg/mg | Estimated | Cell biomass yield coefficient[32] |
| Ratio | 1, 5, 10, 20 | - | Pre-determined | Inoculation Ratio |

**Table S8. Variables and initial conditions in orthogonal obligate commensalism co-culture model**

| Changing variables | Initial Conditions | Units | Constraints | Purpose |
|---|---|---|---|---|
| $X_p$ | 2.5 | mg/L | | Producer biomass |
| $X_u$ | $R*X_p$ | mg/L | | Utilizer biomass |
| $S_f$ | 1000 | mg/L | Sf >= 0 | Feed source concentration |
| N | 0 | mg/L | So >= 0 | OMeTyr concentration |

Supplementary Figures:

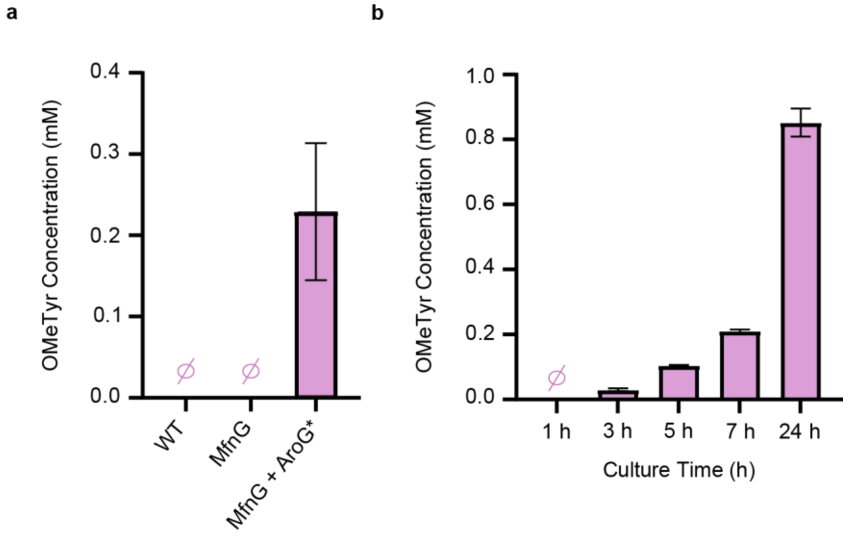

**Supplementary Figure 1.** Optimization of OMeTyr production. **a,** OMeTyr titer values resulting from 24 h growth of RR3 (DE3) *E. coli* without *mfnG* expression (WT), with *mfnG* expression (MfnG), and with co-expression of *mfnG* and *aroG** genes on separate, inducible plasmids (MfnG + AroG*) grown in M9 minimal media supplemented with glucose. Cells were induced at inoculation and grown for 24 h at 34 °C prior to harvesting supernatant samples to check for OMeTyr production. **b,** Production of OMeTyr during a time course carried out in LB media with RR3 (DE3) *E. coli* induced at inoculation. Cells were grown at 34 °C with supernatant samples removed at the timepoints indicated as culture time. Titers were measured through HPLC analysis with peak formations seen using standards as low as 0.01 mM OMeTyr. Sample size is n=3 using biological replicates (triplicate conditions from the same glycerol stock). Data shown are mean ± standard deviation.

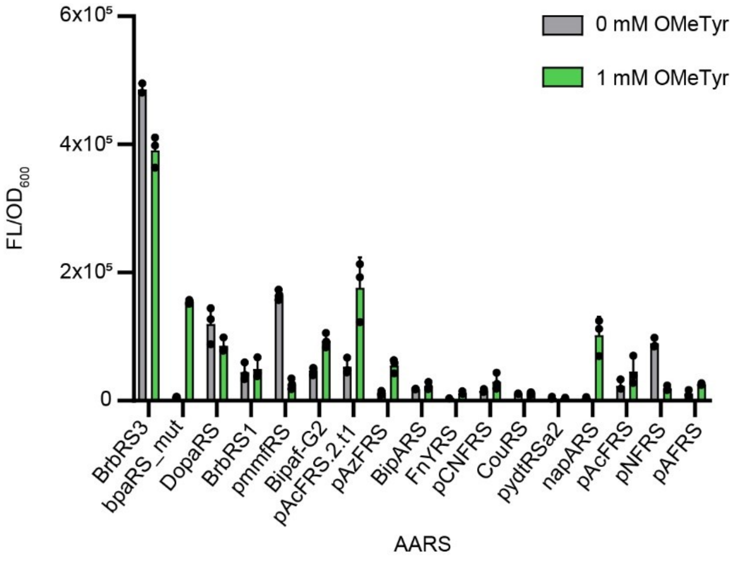

**Supplementary Figure 2.** Initial synthetase screen for incorporation of OMeTyr carried out using various *Methanocaldococcus jannaschii* aminoacyl-tRNA synthetase and tRNA pairs co-expressed with a ubiquitin-fused superfolder GFP. Plasmids were co-expressed in the MG1655 (DE3) strain of *E. coli* in an aromatic amino acid dropout version of Mops EZ Rich media with glucose and 0 or 1 mM OMeTyr supplementation. Cells were grown at 37 °C to mid-exponential phase prior to induction and then returned to 37 °C for 18 h before $OD_{600}$ and fluorescent output (excitation= 485 nm, emission= 525 nm) were measured using a SpectraMax i3x plate reader. Sample size is n=3 using biological replicates. Data shown are mean ± standard deviation.

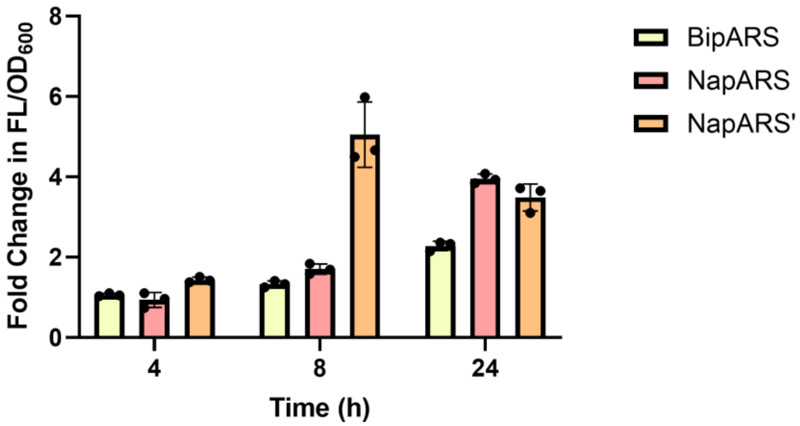

**Supplementary Figure 3.** Fold change in fluorescence (ex: 485 nm, em: 525 nm) normalized by $OD_{600}$ resulting from the incorporation of 0 mM OMeTyr compared to 0.5 mM OMeTyr externally supplied to cultures of the Recoded RARE *E. coli* strain that harbored either the BipARS synthetase, NapARS synthetase, or NapARS' synthetase in LB media. Sample size is n=3 using biological replicates. Data shown are mean ± standard deviation.

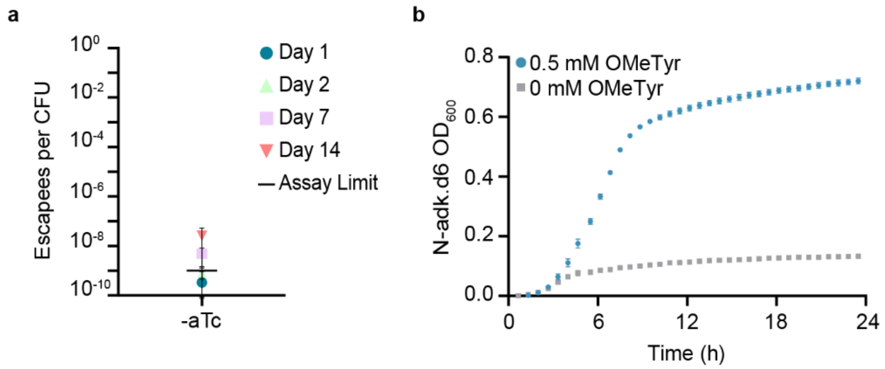

**Supplementary Figure 4.** Escape frequency and growth of N-adk.d6 without aTc induction of synthetase. **a,** Escapees/ CFU calculated as colonies observed on non-permissive media (-aTc, -OMeTyr) per average viable CFU plated for the strain. **b,** Growth comparison of N-adk.d6 grown for 24 h at 34 °C in LB media containing 0.5 mM OMeTyr with and without aTc for synthetase induction. $OD_{600}$ monitoring was carried out using a SpectraMax i3x plate reader. Sample sizes are n=3 using biological replicates and data shown are mean ± standard deviation.

**a**

| Position | 98 | 99 | 100 |
|---|---|---|---|
| | **E** | **A** | **G** |
| A | -58 | n/a | -14 |
| R | -55 | -68 | -17 |
| N | -75 | -76 | -77 |
| D | -74 | -21 | -14 |
| C | -16 | -62 | 11 |
| Q | -89 | -78 | -51 |
| E | n/a | -57 | -16 |
| G | -25 | -40 | n/a |
| H | -25 | -54 | -9 |
| I | -1 | -71 | 36 |
| L | -1 | -79 | -5 |
| K | -67 | -51 | -42 |
| M | -1 | -49 | 29 |
| F | 13 | -46 | 40 |
| P | 32 | -1 | 40 |
| S | -56 | -79 | -33 |
| T | -54 | -64 | -19 |
| W | 26 | -24 | 48 |
| Y | 13 | -57 | 36 |
| V | -42 | -65 | 28 |

**b**

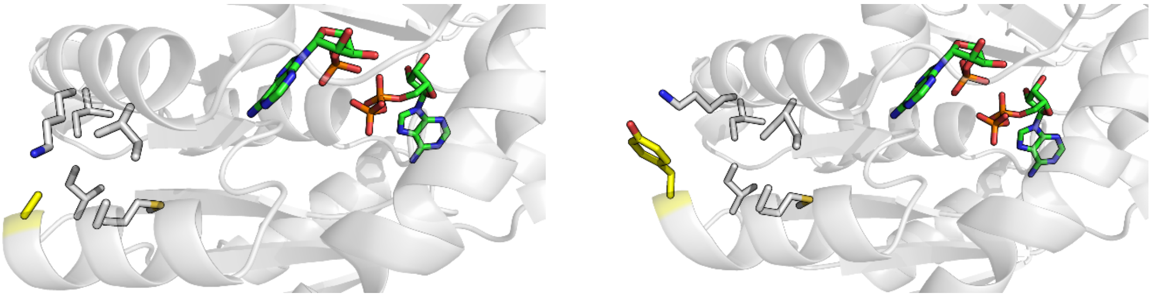

**Supplementary Figure 5.** Sequence conservation and structure-guided analyses used to select position 99 as a candidate for substitution to an nsAA in Adk.d6. **a,** SNAP2 outcome showing the predicted effect of all possible point mutations at position A99 and adjacent amino acids for comparison. A score greater than 50 indicates a strong signal for effect, a score between -50 and 50 indicates a weak signal for effect, and a score less than -50 signals either a neutral or no effect. **b,** (Left) Structural analysis of Adk.d6 with a focus on position A99 (yellow). Relevant residues within 5 Å are shown to provide context for the potential interactions. AMP and ADP (green) binding are also shown using an overlay of the original Adk to demonstrate distance of the intended mutation from the ligand binding pockets. (Right) We perform a very preliminary analysis of how position 99 might be able to accommodate a bulky residue similar to OMeTyr by using PyMOL mutagenesis wizard to substitute A99Y. We then selected relevant proximal residues within 4 Å of the substituted sidechain and performed the "Clean" function, which adjusts the sidechain rotamers to minimize clash.

a

| Position | 250 | 251 | 252 |
| --- | --- | --- | --- |
| | **E** | **Y** | **P** |
| A | -57 | -33 | -50 |
| R | 27 | -46 | -50 |
| N | -58 | -64 | -70 |
| D | -69 | -31 | -62 |
| C | -63 | 3 | -14 |
| Q | -16 | -69 | -76 |
| E | n/a | -43 | -60 |
| G | -11 | -29 | -36 |
| H | -11 | -32 | 0 |
| I | -71 | -11 | -39 |
| L | -88 | -50 | -29 |
| K | 18 | -10 | -73 |
| M | -72 | -28 | -6 |
| F | -75 | -52 | 9 |
| P | 26 | -61 | n/a |
| S | -40 | -59 | -73 |
| T | -78 | -51 | -64 |
| W | -50 | 34 | 1 |
| Y | -76 | n/a | 7 |
| V | -77 | -52 | -49 |

b

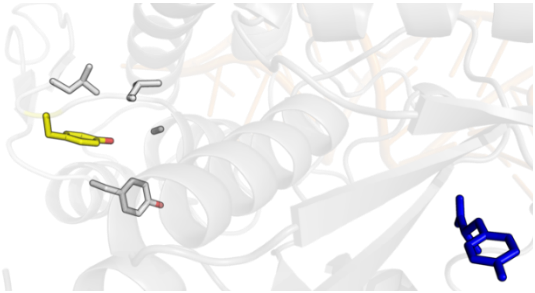

**Supplementary Figure 6.** Rational design considerations applied for UAG codon substitution in NapARS'. **a,** SNAP2 outcome showing the predicted effect of all possible point mutations at position Y251 and adjacent amino acids. **b,** Structural analysis of NapARS' with a focus on position Y251 (yellow). Relevant residues within 5 Å are shown to provide context for the potential interactions. An overlay of the original archaeal tyrosyl-tRNA synthetase complexed with tRNA(Tyr) and L-tyrosine (blue) are also shown to demonstrate distance of the intended mutation from the ligand binding pockets.

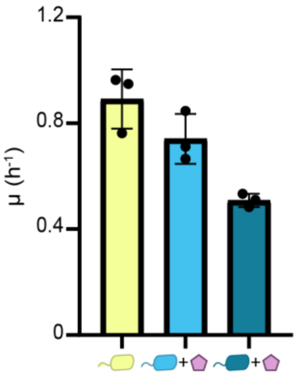

**Supplementary Figure 7.** Specific growth rates calculated for the OMeTyr producing strain (left), N-adk.d6 when supplied 0.5 mM OMeTyr (center), and N.d1-adk.d6 when supplied 0.5 mM OMeTyr (right). Sample sizes are n=3 using biological replicates and data shown are mean ± standard deviation.

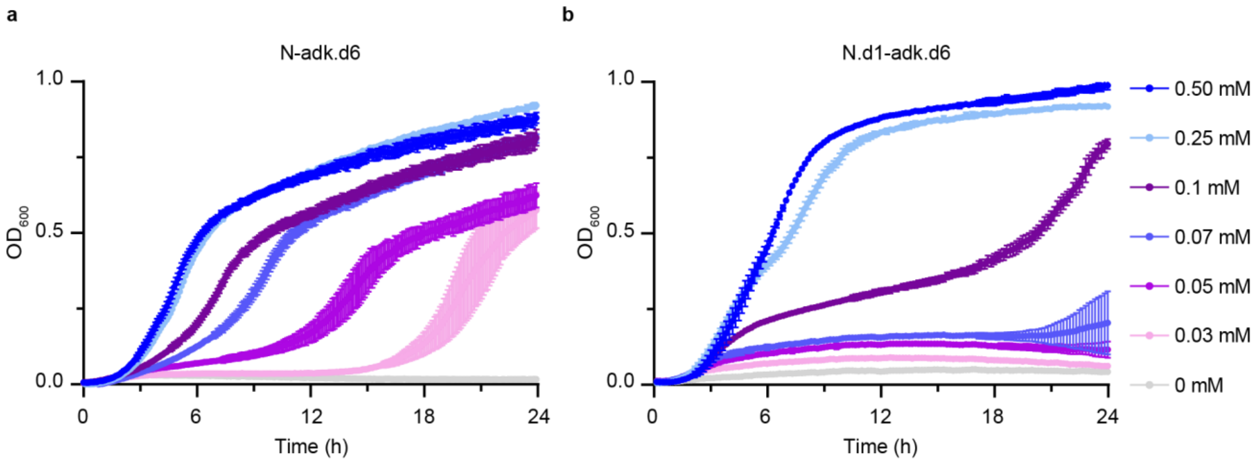

**Supplementary Figure 8.** Growth of synthetic auxotrophs with OMeTyr titration. **a,** N-adk.d6 and **b,** N.d1-adk.d6 growth profiles in LB media for 24 h with decreasing concentrations of externally supplied OMeTyr. $OD_{600}$ monitoring was carried out using a SpectraMax i3x plate reader and a Biotek Synergy H1 plate reader respectively. Sample sizes are n=3 using biological replicates and data shown are mean ± standard deviation.

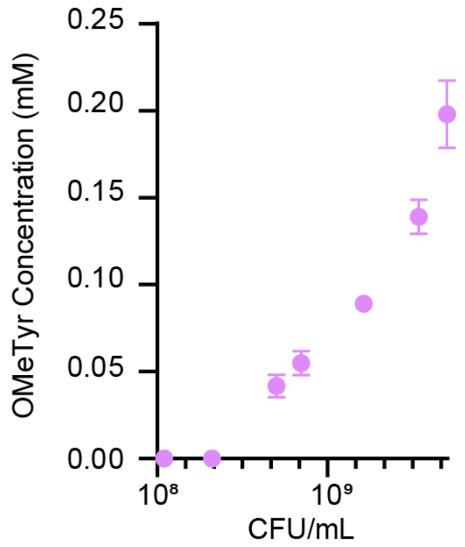

**Supplementary Figure 9.** Titers of OMeTyr production by OMeTyr producer in LB media corresponding to colony counts of the producer strain as CFU/mL. Sample sizes are n=3 using biological replicates and data shown are mean ± standard deviation.

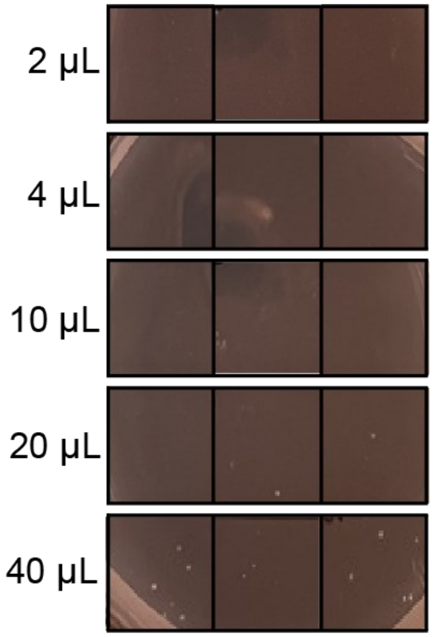

**Supplementary Figure 10.** Persistence of N.d1-adk.d6 as measured by growth on its permissive agar (OMeTyr$^+$) following growth in non-permissive (OMeTyr$^-$) liquid LB media for 24 h at 34 °C. Volume of washed N.d1-adk.d6 used to inoculate liquid culture (left) and images of n=3 replicates on permissive agar plates depicting growth results 24 h after plating on permissive agar (right).

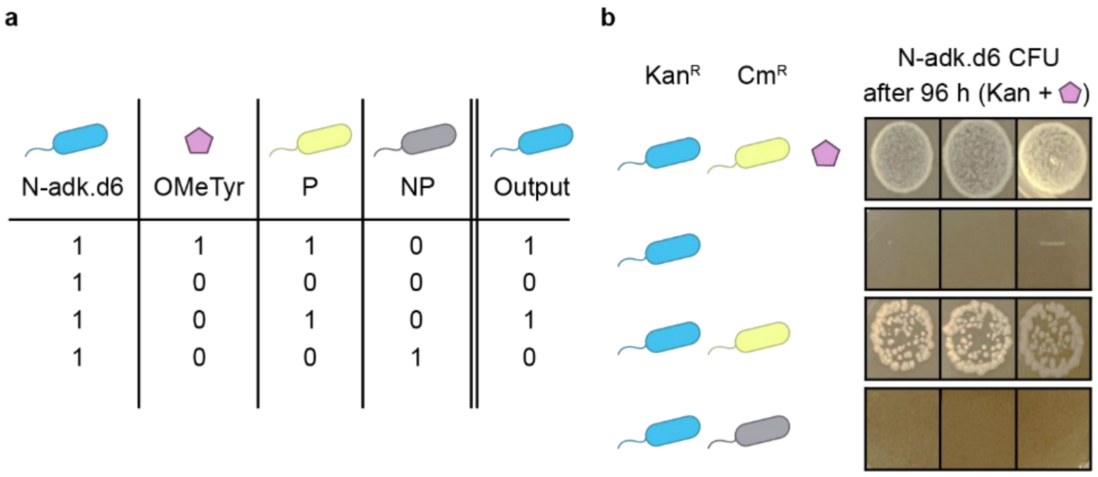

**Supplementary Figure 11.** Obligate commensalism mediated by OMeTyr exemplified using N-adk.d6 synthetic auxotroph. **a,** Predicted outcome of N-adk.d6 survival based on various culture conditions. **b,** Co-culture results depicting n=3 replicates for each condition listed. 1st row: N-adk.d6 co-cultured with OMeTyr producer at a 1:1 inoculation ratio grown in permissive media containing 0.5 mM externally supplied OMeTyr (pentagon). 2nd row: N-adk.d6 monoculture grown in non-permissive media. 3rd row: N-adk.d6 co-cultured with OMeTyr producer at a 1:1 inoculation ratio in non-permissive media. 4th row: N-adk.d6 co-cultured with a non-producing *E. coli* strain at a 1:1 inoculation ratio in non-permissive media. Images shown represent colonies 96 h after plating on solid media selective for N-adk.d6.

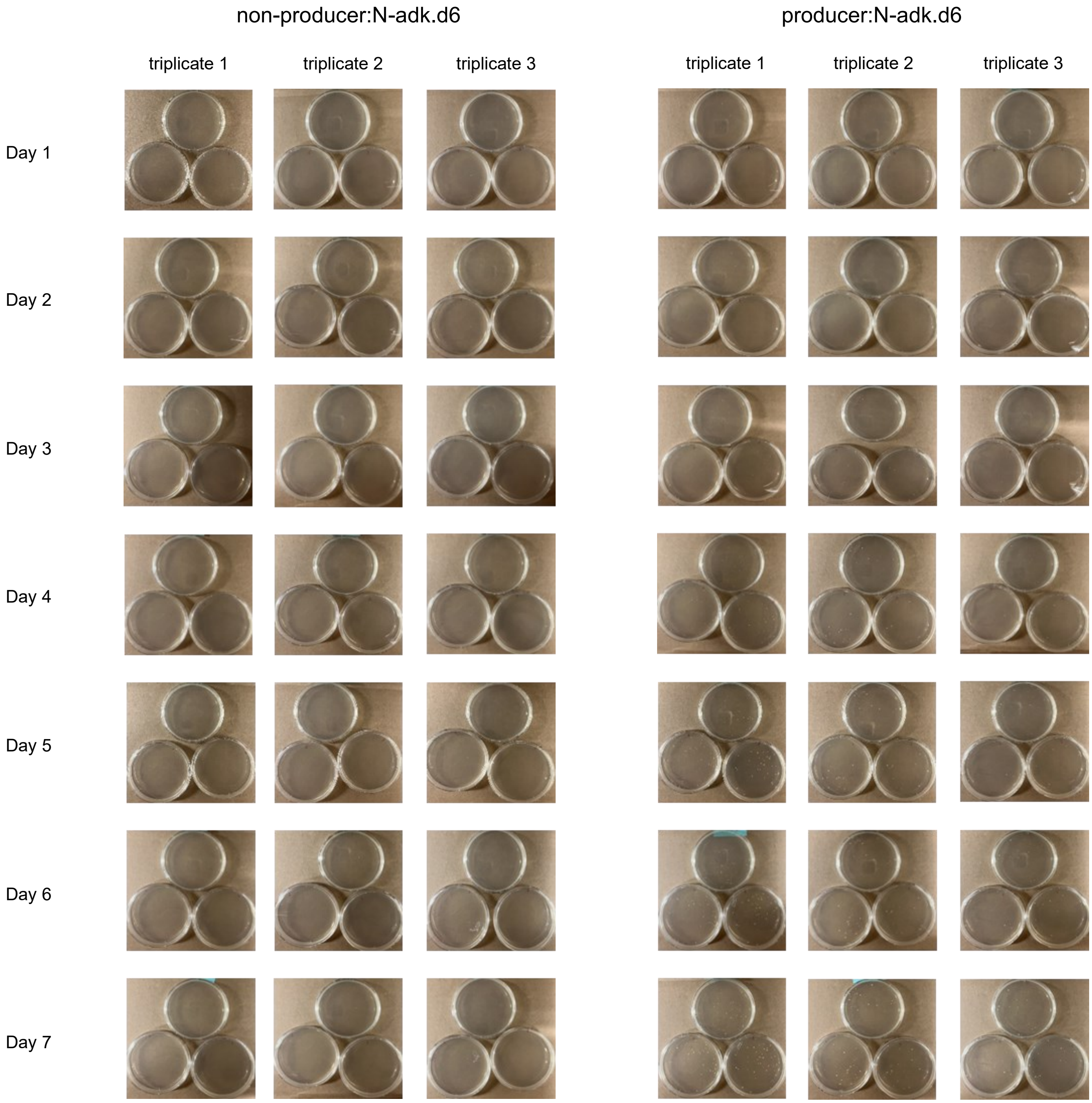

**Supplementary Figure 12.** Growth of full co-culture volumes on non-permissive agar selective for N-adk.d6. Each of the triplicate wells from the 1:1 (non-)producer:N-adk.d6 co-cultures were washed in LB and the entirety of the culture volumes were plated across 3 non-permissive agar plates with kanamycin to select for N-adk.d6 growth. Growth was monitored for 7 days at 34 °C.

a

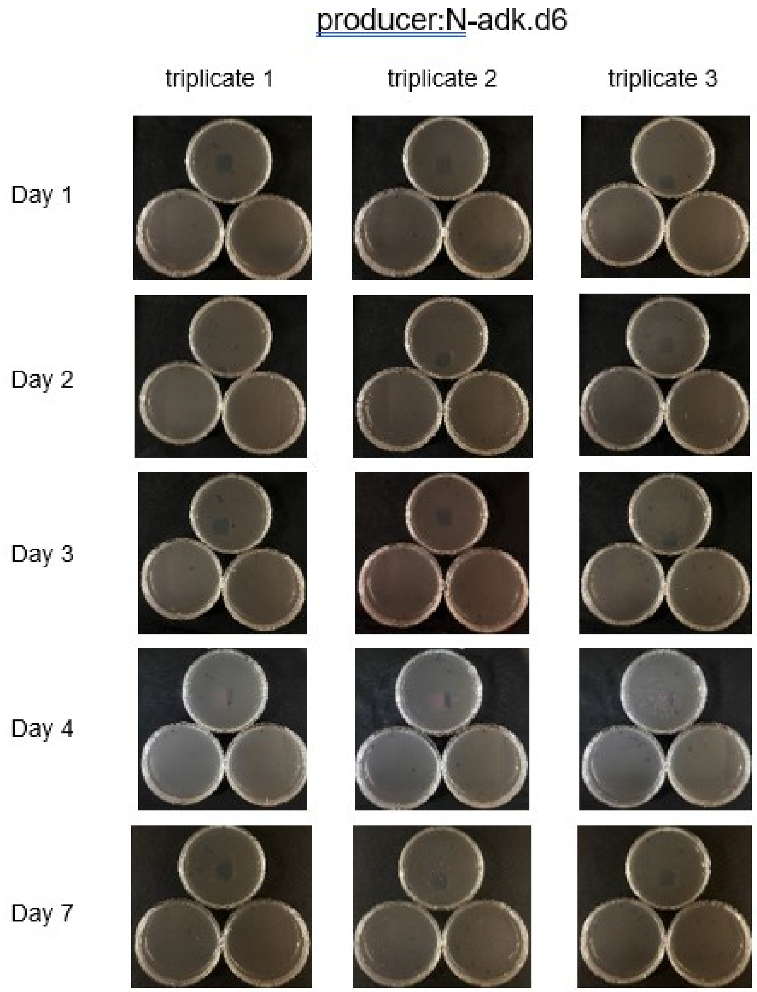

b

| | | | | | | | | | | | |
|---|---|---|---|---|---|---|---|---|---|---|---|
| 0.008933 | 0.003333 | 0.004733 | 0.004933 | 0.008533 | 0.006533 | 0.007133 | 0.008733 | 0.005333 | 0.003533 | 0.057133 | 0.006733 |
| 0.007333 | 0.007533 | 0.005533 | 0.007933 | 0.130133 | 0.008533 | 0.007133 | 0.006933 | 0.014933 | 0.004933 | 0.006733 | |
| 0.166733 | 0.006333 | 0.245933 | 0.007333 | 0.046933 | 0.634733 | 0.132133 | 0.014733 | 0.011933 | 0.486533 | 0.007133 | 0.007333 |
| 0.011333 | 0.008533 | 0.719733 | 0.011933 | 0.014933 | 0.019333 | 0.168533 | 0.011933 | 0.009333 | 0.502733 | 0.740933 | 0.006533 |
| 0.059333 | 0.008933 | 0.009133 | 0.011333 | 0.024533 | 0.004333 | 0.039733 | 0.011133 | 0.010933 | 0.189933 | 0.015933 | 0.007933 |
| 0.021133 | 0.007533 | 0.047133 | 0.013333 | 0.016733 | 0.005733 | 0.006933 | 0.010133 | 0.008733 | 0.020333 | 0.005933 | 0.019333 |
| 0.054133 | 0.006733 | 0.033333 | 0.357533 | 0.018533 | 0.015133 | 0.009533 | 0.203333 | 0.101133 | 0.018533 | 0.008133 | 0.006533 |
| 0.1672 | 0.0018 | 0.0022 | 0.0038 | | | | | | | | |
| 0.0922 | 0.003 | 0.006 | 0.2424 | 0.0244 | 0.0208 | 0.005 | 0.0148 | 0.0042 | 0.0042 | 0.1352 | |

**Supplementary Figure 13.** Escapee monitoring from full co-culture volume plated on non-permissive agar selective for N.d1-adk.d6 following 1:5 producer:N.d1-adk.d6 co-cultures. **a,** Each replicate was washed in LB and the entirety of the culture volumes were plated across 3 non-permissive agar plates with kanamycin to select for N.d1-adk.d6 growth. Growth was monitored for 7 days at 34 °C. **b,** Following the 7-day incubation on solid media, each colony that appeared across the replicate plates was picked and grown for 72 h in non-permissive LB liquid media (OMeTyr$^{-}$) at 34 °C. A SpectraMax i3x plate reader was used to determine $OD_{600}$ following liquid outgrowth. Individual $OD_{600}$ values are shown for each colony. Colonies that resulted in outgrowth to at least 0.1 $OD_{600}$ after 72 h were deemed viable escapees.

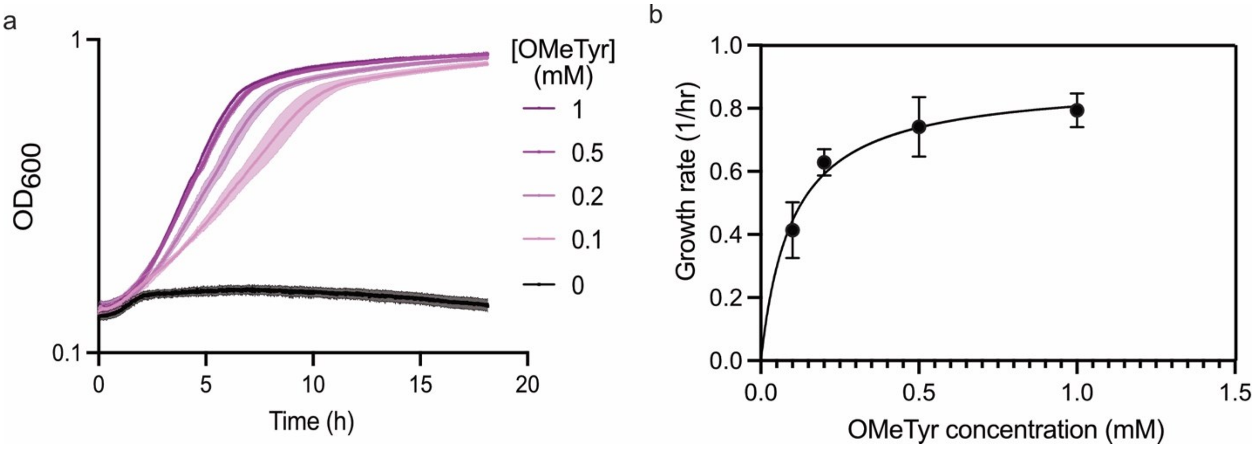

**Supplementary Figure 14** Parameter estimation for co-culture model. **a,** Growth curves for the utilizer at different concentrations of OMeTyr. The line represents the mean of biological triplicates with the standard deviation represented as the color shading around the line. **b,** Growth rate versus OMeTyr concentration to estimate the Monod growth kinetics parameters. The solid line represents a non-linear regression performed by GraphPad Prism v9. Growth rate data was calculated by performing a linear regression on the linear region of the growth curves in **a**.

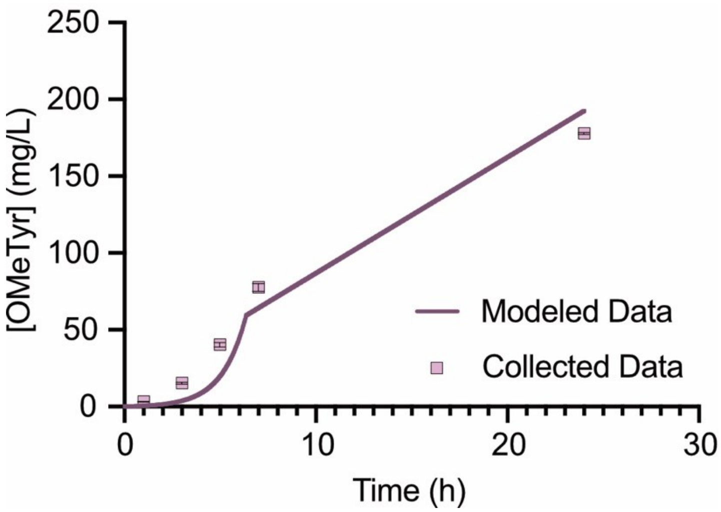

**Supplementary Figure 15.** Model validation by comparing the modeled concentration of OMeTyr produced over time compared to the actual concentration of OMeTyr produced as shown in Fig 1d.

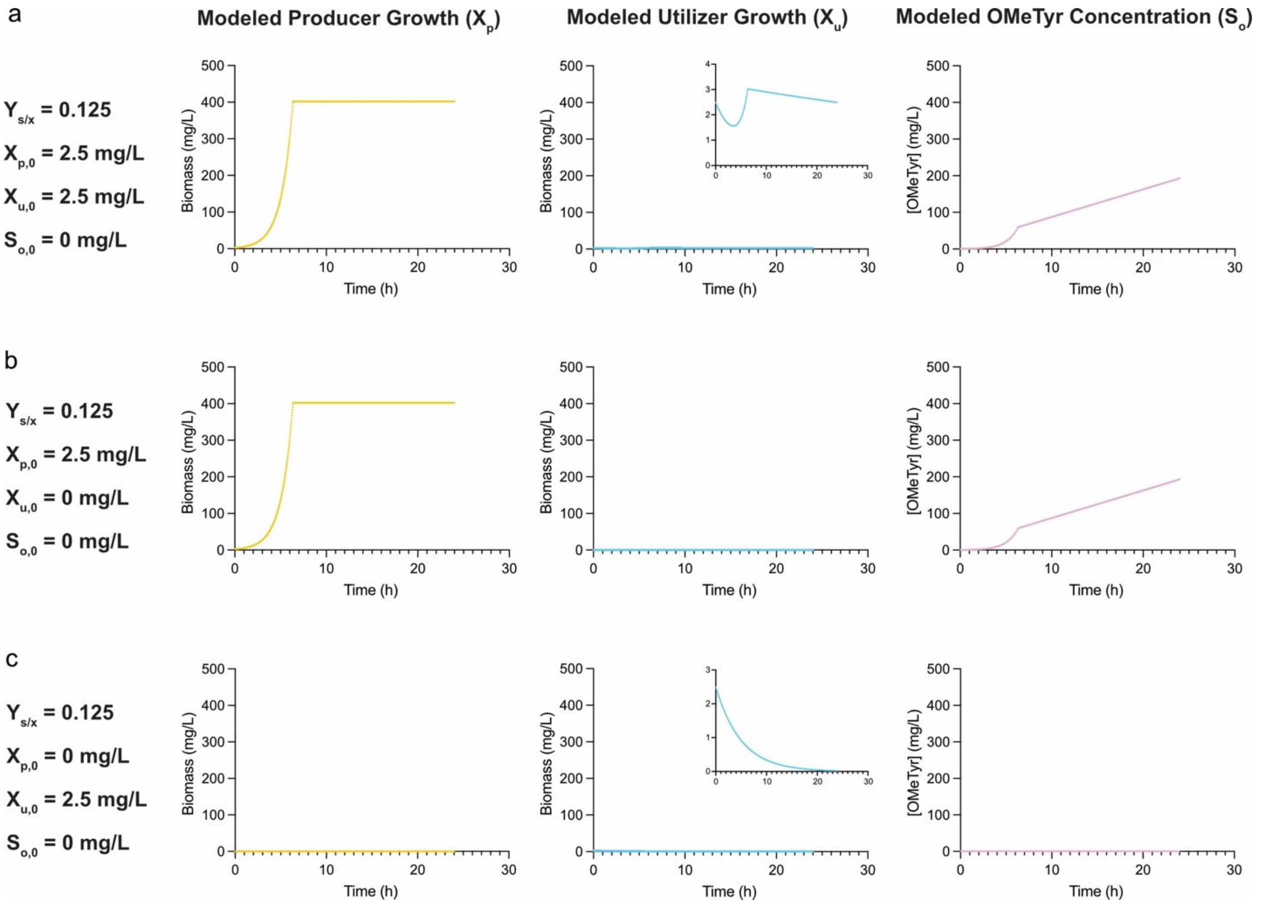

**Supplementary Figure 16.** Model controls to confirm the predicted growth of the producer and utilizer (N-adk.d6) is consistent with expectations in limiting cases. We included insets where there was non-constant behavior for the culture that was not visible with the original the axis scaling, otherwise the behavior was constant. **a**, The predicted behavior for the producer, utilizer, and OMeTyr concentration in the model if the utilizer and producer are inoculated in a 1:1 ratio. **b**, Modeled behavior for when the producer is in a monoculture. **c**, Modeled behavior for when the utilizer is in a monoculture.

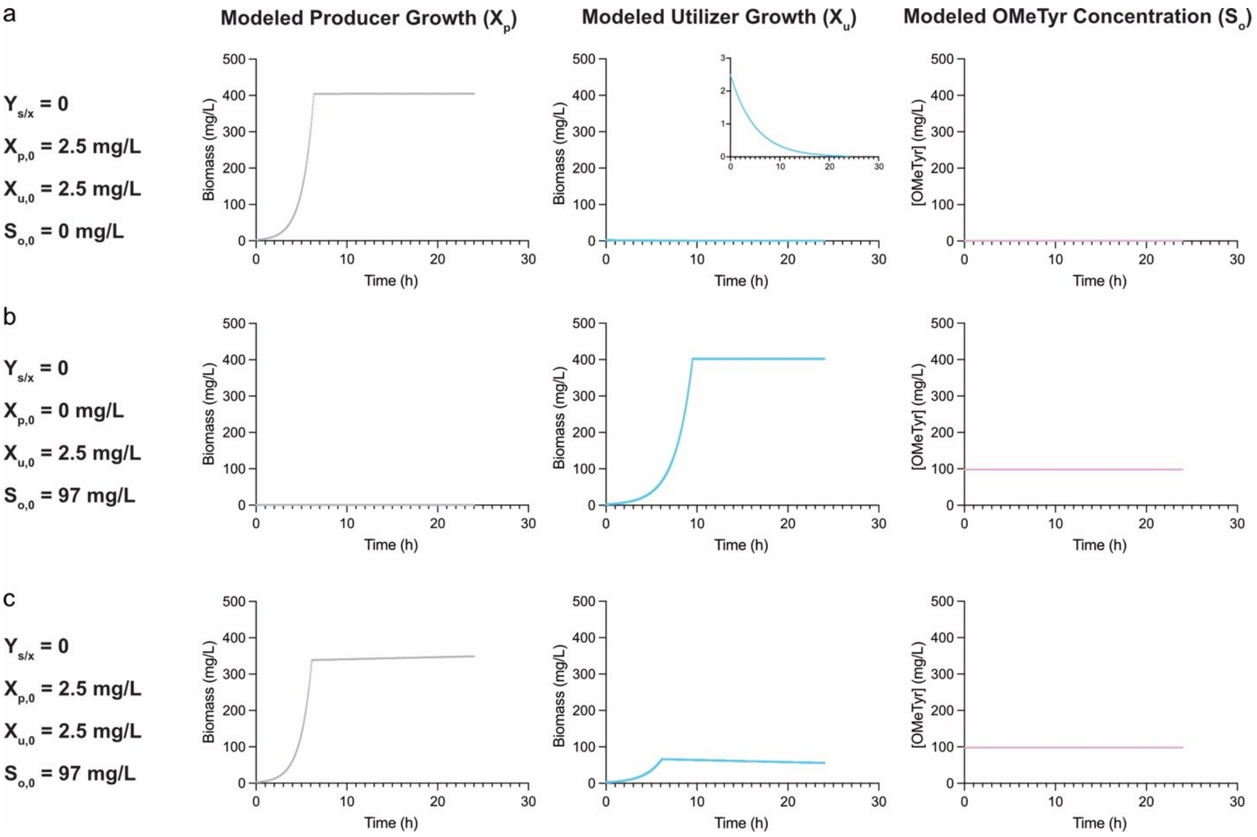

**Supplementary Figure 17.** Model controls to confirm the predicted growth of the non-producer and utilizer (N-adk.d6) is consistent with expectations in limiting cases including in the presence of supplemented OMeTyr. We included insets where there was interesting behavior for the culture that was not visible with the original the axis scaling, otherwise the behavior was constant. **a**, Model behavior when the non-producer is present in a co-culture with the utilizer. **b**, Model behavior when the utilizer is present in a monoculture with supplemented OMeTyr. **c**, When OMeTyr is supplemented but the non-producer is present, there is a balance between the non-producer and utilizer.

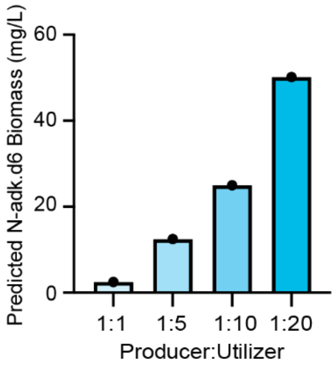

**Supplementary Figure 18.** Simulated final cell densities as a function of inoculation ratio based on kinetic growth modeling.

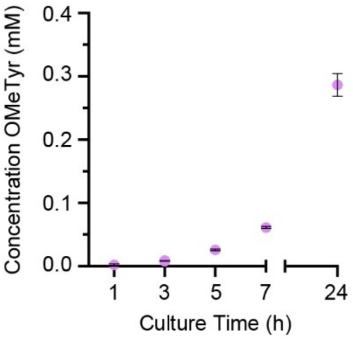

**Supplementary Figure 19.** Titers of OMeTyr as a function of time produced by DEP.e5 that co-expresses *mfnG* and *aroG** genes under a constitutive promoter in its permissive LB media containing 10 µM BipA. Sample sizes are n=3 using biological replicates and data shown are mean ± standard deviation.

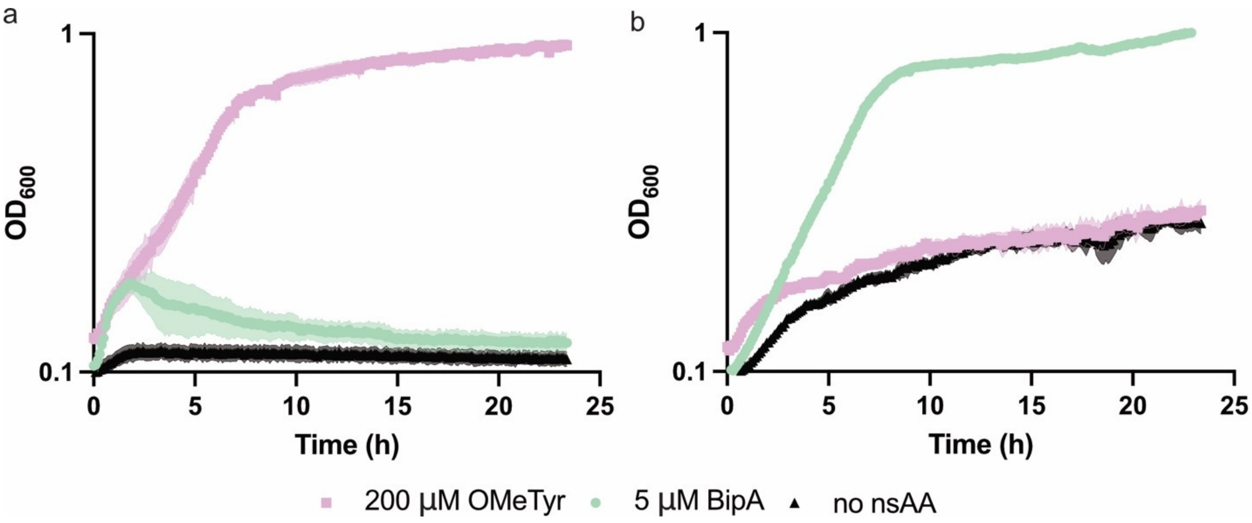

**Supplementary Figure 20.** Growth curves of **a**, N-adk.d6 and **b**, DEP.e5 with different nsAAs. N-adk.d6 can only grow on OMeTyr demonstrated by the robust growth with OMeTyr and rapid death with no nsAA and BipA. DEP.e5 appears to only grow on BipA as demonstrated by the alignment of the low-density curves for DEP.e5 with OMeTyr and no nsAA. Some initial outgrowth and persistence after removal of BipA has been observed and reported previously for DEP and its descendant strains.

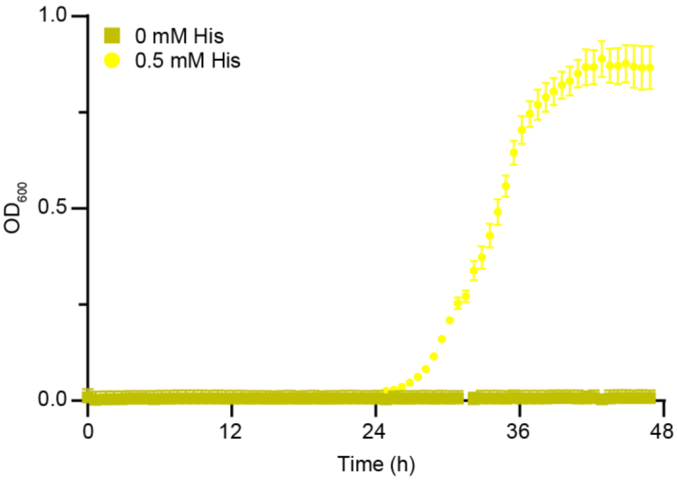

**Supplementary Figure 21.** Kinetic growth profiles of Δ*hisD* producer strain in M9 minimal media with and without the external addition of 0.5 mM histidine. $OD_{600}$ monitoring was carried out using a SpectraMax i3x plate reader. Sample sizes are n=3 using biological replicates and data shown are mean ± standard deviation.

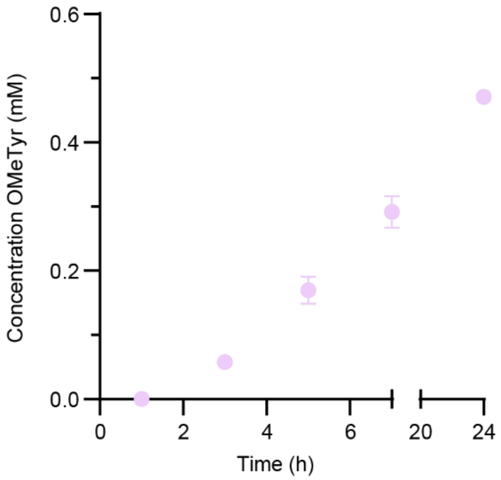

**Supplementary Figure 22.** Titers of OMeTyr as a function of time produced by Δ*hisD* that co-expresses *mfnG* and *aroG** genes under a constitutive promoter in LB media. Sample sizes are n=3 using biological replicates and data shown are mean ± standard deviation.

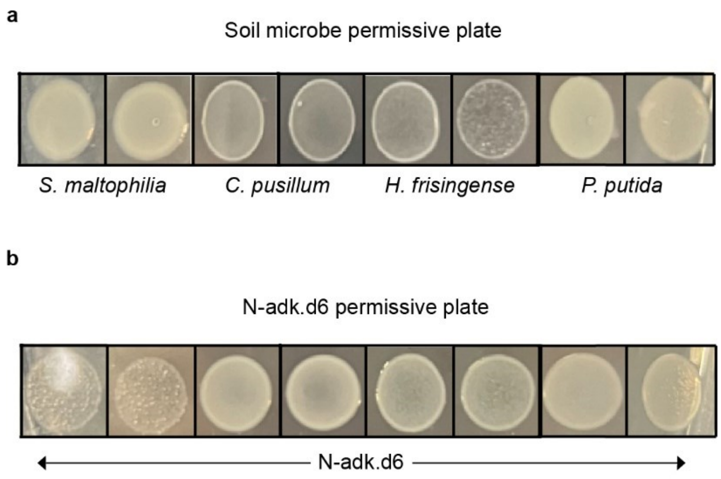

**Supplementary Figure 23.** Growth of individual organisms on permissive plates following 24 h co-culture in N-adk.d6 permissive LB media containing 0.5 mM OMeTyr. **a.** Growth of individual soil microbes following co-culture with N-adk.d6 in the synthetic auxotroph's permissive media without antibiotics. Soil microbes plated on LB agar, which is non-permissive for the growth of N-adk.d6. **b.** Growth of N-adk.d6 on its own permissive agar following co-culture with *S. maltophilia, C. pusillum, H. frisingense,* or *P. putida*. Agar contains 30 µg/mL kanamycin, which selects for the growth of N-adk.d6 but not the growth of the soil microbes. Sample size is n=2 using biological replicates plated separately.

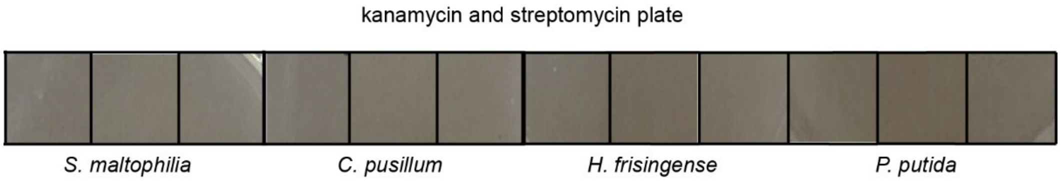

**Supplementary Figure 24.** Growth investigation of individual soil microbes on agar plates containing 15 μg/mL kanamycin and 47.5 streptomycin μg/mL, which selects for the growth of N-adk.d6 but not the growth of the soil microbes. Plate stored at 34 °C for 24 h. Sample size is n=3 using biological replicates plated separately.

Non-producer:soil consortium:N.d1-adk.d6-strep

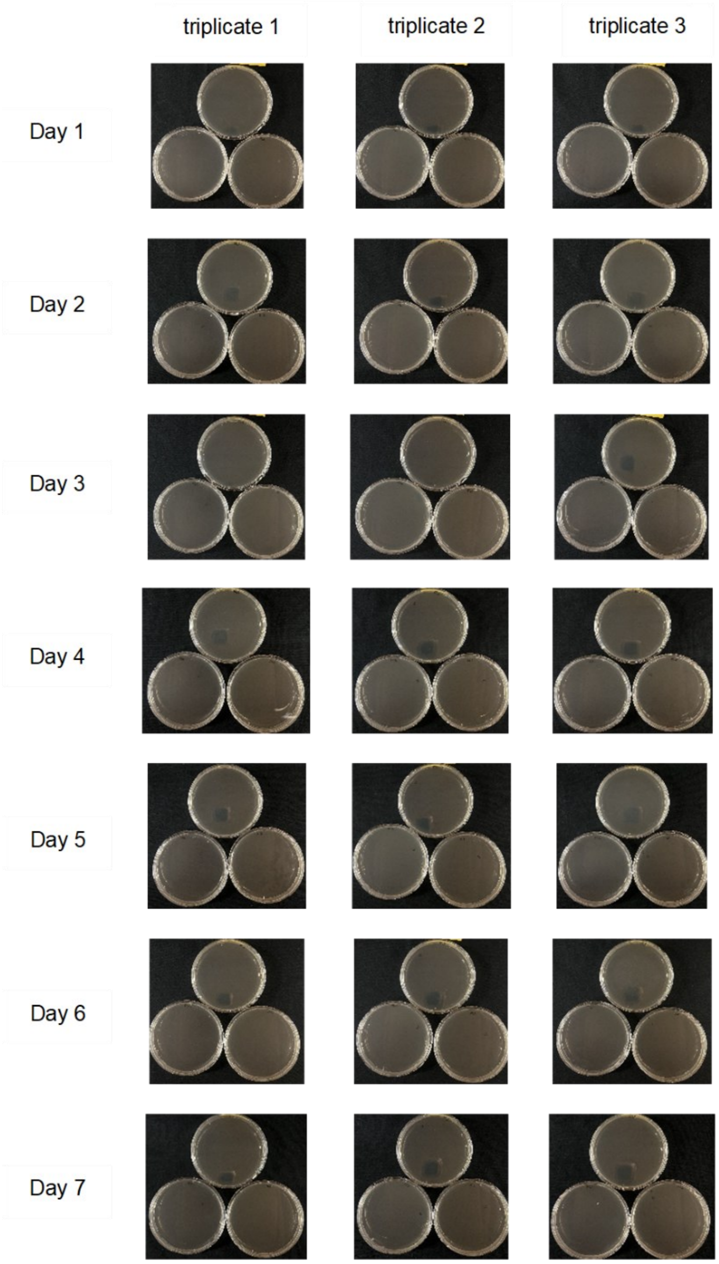

**Supplementary Figure 25.** Growth of full co-culture volumes on non-permissive agar selective for N.d1-adk.d6-strep. Each of the triplicate wells from the 1:1:10 non-producer:soil consortium:N.d1-adk.d6-strep co-cultures were washed in LB and the entirety of the culture volumes were plated across 3 non-permissive agar plates with kanamycin and streptomycin to select for N.d1-adk.d6-strep growth. Growth was monitored for 7 days at 34 °C.
